# Dentate Gyrus and CA3 GABAergic Interneurons Bidirectionally Modulate Signatures of Internal and External Drive to CA1

**DOI:** 10.1101/2021.01.04.425303

**Authors:** Emily A. Aery Jones, Antara Rao, Misha Zilberter, Biljana Djukic, Anna K. Gillespie, Nicole Koutsodendris, Maxine Nelson, Seo Yeon Yoon, Kylie Huang, Heidi Yuan, Theodore M. Gill, Yadong Huang, Loren M. Frank

**Author notes:** Correspondence should be addressed to: Loren Frank or Yadong Huang.

## Abstract

Specific classes of GABAergic neurons are thought to play specific roles in regulating information processing in the brain. In the hippocampus, two major classes – parvalbumin-expressing (PV^+^) and somatostatin-expressing (SST^+^) neurons – differentially regulate endogenous firing patterns and target different subcellular compartments of principal cells, but how these classes regulate the flow of information throughout the hippocampus is poorly understood. We hypothesized that PV^+^ and SST^+^ interneurons in the dentate gyrus (DG) and CA3 might differentially modulate CA3 patterns of output, thereby altering the influence of CA3 on CA1. We found that while suppressing either interneuron type increased DG and CA3 output, the effects on CA1 were very different. Suppressing PV^+^ interneurons increased local field potential signatures of coupling from CA3 to CA1 and decreased signatures of coupling from entorhinal cortex to CA1; suppressing SST^+^ interneurons had the opposite effect. Thus, DG and CA3 PV^+^ and SST^+^ interneurons bidirectionally modulate the flow of information through the hippocampal circuit.

## INTRODUCTION

GABAergic interneurons regulate principal cell input and output and act as hubs for controlling network activity throughout the brain (McKenzie, 2017). These interneurons are highly heterogeneous (Klausberger and Somogyi, 2008), and distinct classes may play distinct roles in balancing internal and external input drive to local circuits.

In the hippocampus, a brain structure critical for encoding, consolidation, and retrieval of memories, the majority of GABAergic interneurons are either parvalbumin-expressing (PV^+^) or somatostatin-expressing (SST^+^) (Jinno and Kosaka, 2002, 2003). PV^+^ and SST^+^ interneurons throughout the hippocampus are distinguished by their subcellular spatial domains, firing properties, pyramidal cell spike modulation mechanisms, and temporal coordination (Klausberger and Somogyi, 2008). These two interneuron classes are well characterized in CA1, where they are uniquely positioned to bidirectionally regulate information flow from the two major inputs: internal input from CA3 and external input from entorhinal cortex (EC). PV^+^ interneurons mainly target somatic and perisomatic compartments and thus regulate principal cell excitability and precise spike timing (Klausberger and Somogyi, 2008; Lovett-Barron et al., 2012; Miles et al., 1996; Royer et al., 2012). In CA1, these cells are preferentially recruited by excitatory EC inputs and are thus the main drivers of feedforward inhibition (Freund and Buzsáki, 1998; Gulyás et al., 1999; Lee et al., 2016; Wheeler et al., 2015). In contrast, SST^+^ interneurons mainly target distal apical dendrites – which receive extrahippocampal excitatory inputs from the EC and medial septum (MS) – and gate the influence of those pathways, thereby affecting the magnitude of principal cell spiking (Blasco-Ibáñez and Freund, 1995; Klausberger and Somogyi, 2008; Lovett-Barron et al., 2012; Sik et al., 1995, 1997; Takács et al., 2012). In CA1, these cells receive primarily local inputs from pyramidal cells and thus provide feedback inhibition (Freund and Buzsáki, 1998; Wheeler et al., 2015). Consistent with these patterns of inputs, potentiation of PV^+^ interneurons attenuates CA3 inputs while potentiation of CA1 SST^+^ interneurons attenuates EC inputs (Fernández-Ruiz et al., 2017; Udakis et al., 2020).

Whether PV^+^ and SST^+^ interneurons in the upstream regions of the dentate gyrus (DG) and CA3 play similar distinct roles is unknown. Since both populations are inhibitory, we would expect that suppressing either PV^+^ or SST^+^ interneurons in DG and CA3 would lead to greater total spiking output from CA3 and thus greater input to CA1. Moreover, there is some overlap between PV and SST expression (Jinno and Kosaka, 2000), making it unclear whether broadly manipulating these genetically-defined cell classes would modulate outputs in opposing directions.

However, given their similarities to analogous populations in CA1 (Klausberger and Somogyi, 2008), one can hypothesize that DG and CA3 PV^+^ interneurons provide feed-forward inhibition at the soma, thus suppressing internal DG to CA3 and CA3 to CA3 inputs. Conversely, as DG and CA3 SST^+^ interneurons are thought to provide feedback inhibition at the level of EC inputs, we would expect them to suppress those external inputs. Thus, suppressing DG and CA3 PV^+^ interneurons might ungate internal drive in DG and CA3, while suppressing DG and CA3 SST^+^ interneurons might ungate external drive. If the resulting patterns of CA3 activity differentially engage CA1, we would predict that suppressing either PV^+^ or SST^+^ interneurons in DG and CA3 would enhance DG and CA3 spiking outputs, but this increase in output could have different effects on CA1.

To measure these potential effects we took advantage of known signatures of information flow in the hippocampal network. During immobility, high CA3 drive to CA1 is associated with sharp-wave ripple (SWR) events (Buzsáki et al., 1992; Csicsvari et al., 2000; Ylinen et al., 1995) that can be detected as increases in ripple frequency (125–200 Hz) power in CA1. Previous work has shown that blocking CA3 input to CA1 reduces SWR rate, slows SWR frequency, and increases multi-unit activity (MUA) recruitment to SWRs (Middleton and McHugh, 2016; Nakashiba et al., 2009; Yamamoto and Tonegawa, 2017) while increasing CA3 drive to CA1 increases SWR rate, SWR frequency, SWR-coincident slow gamma (SG) power, and sharp wave (SW) amplitude (Ramirez-Villegas et al., 2018; Wu et al., 2015). By contrast, blocking direct EC input to CA1 diminishes SWR chains, but has no other observable effects on SWR rate, frequency, or duration (Yamamoto and Tonegawa, 2017), while lesioning EC increases SWR rate (Bragin et al., 1995a). Thus, SWR properties provide information about CA3 and EC drive to CA1, and these signatures are bidirectional: decreased internal CA3 drive is associated with decreased SWR activity, and decreased external EC drive is associated with increased SWR activity.

During movement, EC drive to CA1 is associated with fast gamma (FG; 50–100 Hz) oscillations, while CA3 drive to CA1 is associated with SG (20–50 Hz) oscillations (Bragin et al., 1995b; Colgin, 2016; Colgin et al., 2009). Blocking synaptic inputs from CA3 to CA1 impacts CA1 SG power and modulation by theta phase while not detectably affecting CA1 SG frequency and coherence across regions (Middleton and McHugh, 2016). Direct EC layer III inputs to CA1 have been shown to contribute to CA1 FG activity (Yamamoto et al., 2014). However, lesioning inputs from CA3 to CA1 also reduces CA1 FG power, suggesting that FG may be both internally and externally generated (Middleton and McHugh, 2016). SG and FG are in turn organized by theta (5–11 Hz), a signature of EC and MS drive to CA1 (Colgin, 2015). Theta is not detectably disrupted by blocking CA3 inputs to CA1 (Middleton and McHugh, 2016) but does diminish in power and frequency throughout the hippocampus when EC is lesioned (Bragin et al., 1995b; Buzsáki et al., 1983; Maurer et al., 2006; Ormond et al., 2015; Schlesiger et al., 2015). Thus, theta and gamma rhythms during movement can provide information about internal CA3 and external EC drive to CA1.

We therefore asked whether modulation of specific interneuron types in DG and CA3 could alter these signatures of EC and CA3 drive to CA1. We found that while suppression of either interneuron type led to an increase in DG and CA3 spiking output, suppressing PV^+^ interneurons increased signatures of CA3 coupling to CA1 and decreased those of EC coupling to CA1, while suppressing SST^+^ interneurons decreased signatures of CA3 coupling to CA1 and increased those of EC coupling to CA1. Our findings indicate that PV^+^ and SST^+^ interneurons in DG and CA3 can bidirectionally alter patterns of activity in DG and CA3, altering the balance of LFP signatures of EC and CA3 drive in CA1.

## RESULTS

### Recording *in vivo* hippocampal LFP during chemogenetic suppression of PV^+^ and SST^+^ interneurons in DG and CA3

We used a Cre-dependent chemogenetic approach to silence specific interneuron classes in DG and CA3. We bilaterally injected PV-Cre, SST-Cre, and PV-Cre/SST-Cre mice with AAV5-hSyn-DIO-hM4D(Gi)-mCherry or with AAV5-hSyn-DIO-mCherry into the DG hilus, and then implanted a 32-site silicon electrode array into the right dorsal hippocampus (Figure 1A). HM4D expression was highly co-localized with PV and SST expression (Figure S1A and S1B) and extended throughout DG and CA3 (Figure 1B and S1C) from dorsal to ventral hippocampus, with no expression in CA1 (Figure 1C). We confirmed hM4D function in *ex vivo* brain slices by observing that clozapine N-oxide (CNO) application reduced firing rates in PV^+^ and SST^+^ interneurons expressing hM4D (Figure S1D), hyperpolarized resting membrane potentials by 8.6 ± 1.3mV (Figure S1E), and decreased input resistance by 31.3 ± 9.9 MΩ (Figure S1F). After viral expression, we recorded LFP activity from CA1, CA3, and DG over 6 daily sessions, alternating CNO and vehicle treatment across days (Figure 1A). CNO was used to suppress interneuron activity for the duration of the recording. We collected data from two cohorts of animals: the first underwent only home cage recordings, and the second underwent linear track recordings followed by home cage recordings.

**Figure 1.**
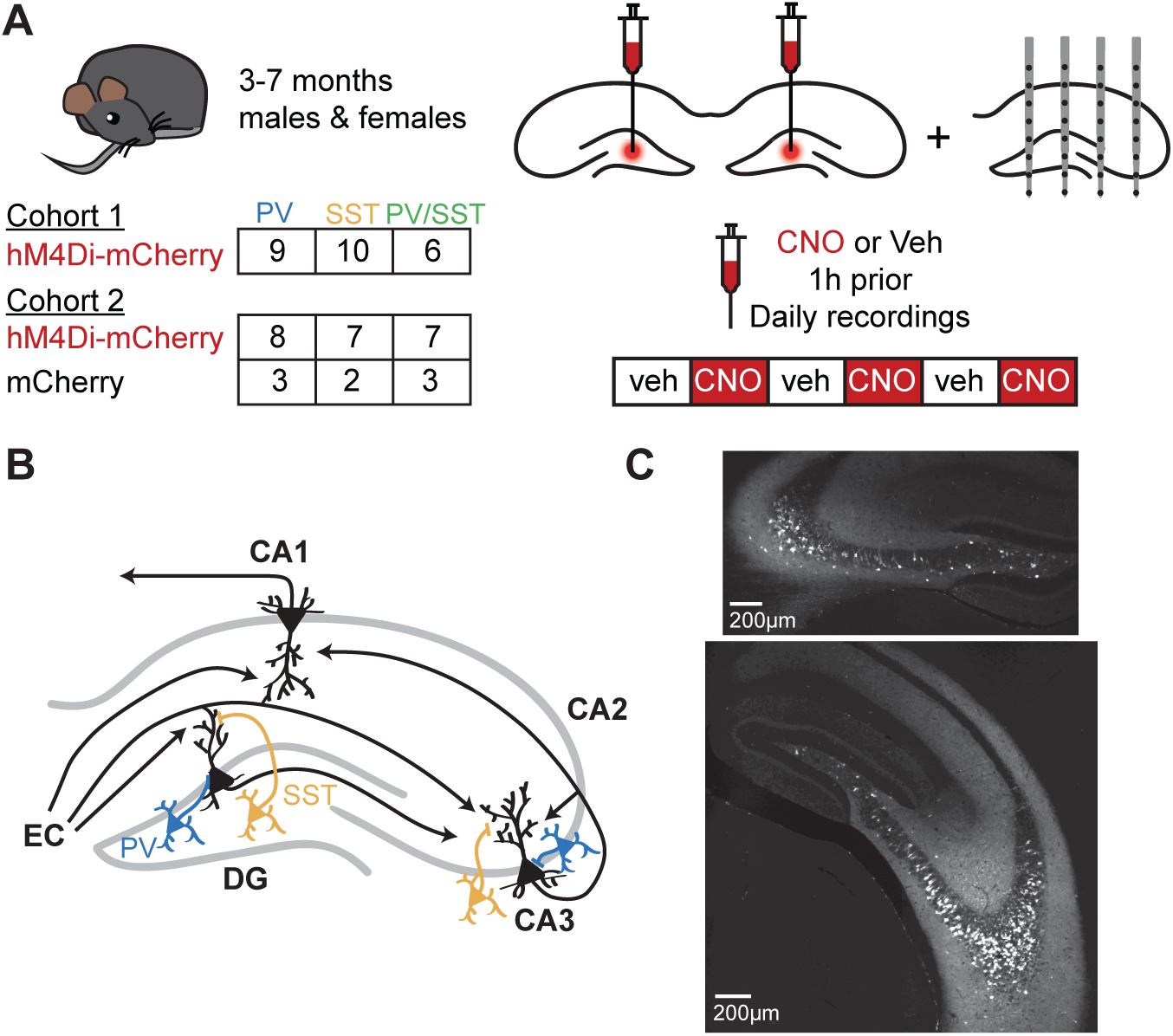
**Recording *in vivo* hippocampal LFP during chemogenetic suppression of PV^+^ and SST^+^ interneurons in DG and CA3.** (A) Experimental strategy. Mice were bilaterally injected with AAV5-hSyn-DIO-hM4D(Gi)- mCherry or with AAV5-hSyn-DIO-mCherry into the DG hilus, then implanted with a 32-site silicon electrode array into the right dorsal hippocampus. LFP activity was recorded from all hippocampal subregions over 6 daily 1 hour sessions, alternating CNO and vehicle treatment. (B) Simplified circuit diagram of the hippocampus. DG and CA3 receive excitatory inputs from EC layer II onto distal dendrites and inhibitory inputs from local PV^+^ interneurons (blue) onto the soma and local SST^+^ interneurons (yellow) onto distal dendrites. CA1 receives direct input from EC layer III, indirect input from EC layer II via DG and CA3, and input from CA3. Adapted with permission from (Gillespie et al., 2016). (C) Example of mCherry expression in dorsal (top) and ventral (bottom) hippocampus. See also Figure S1.

We assessed LFP features as a readout for changes in internal CA3 versus external EC drive to CA1, focusing on features previously identified to be modulated by CA3 or EC drive. We took advantage of a statistical approach known as a Linear Mixed Model (LMM) to assess differences between groups while accounting for variability both within and across individuals. We evaluated differences between significant treatment effects to determine which effects were significantly different between genotypes (referred to as specific). We further sought to control our false positive rate and so only report findings which survive Holm-Bonferroni correction across all comparisons in each experiment (see Methods).

### Suppressing PV^+^ or SST^+^ interneurons bidirectionally modulates SWR signatures of CA3 coupling to CA1

We first asked whether PV^+^ or SST^+^ interneurons in DG and CA3 modulate signatures of CA3 coupling to CA1 as measured through increased SWR activity during immobility (<1 cm/s for ≥30 s) in 1-hour daily home cage sessions (Figure 2A). We hypothesized that suppressing DG and CA3 PV^+^ interneurons might ungate internal DG and CA3 inputs and thus changing DG and CA3 firing patterns in a manner that facilitates internally driven patterns of activity (e.g. SWRs) in CA1. Conversely, suppressing DG and CA3 SST^+^ interneurons might ungate EC inputs and perhaps change DG and CA3 firing patterns in a manner that increases CA1 receptivity to EC drive.

**Figure 2.**
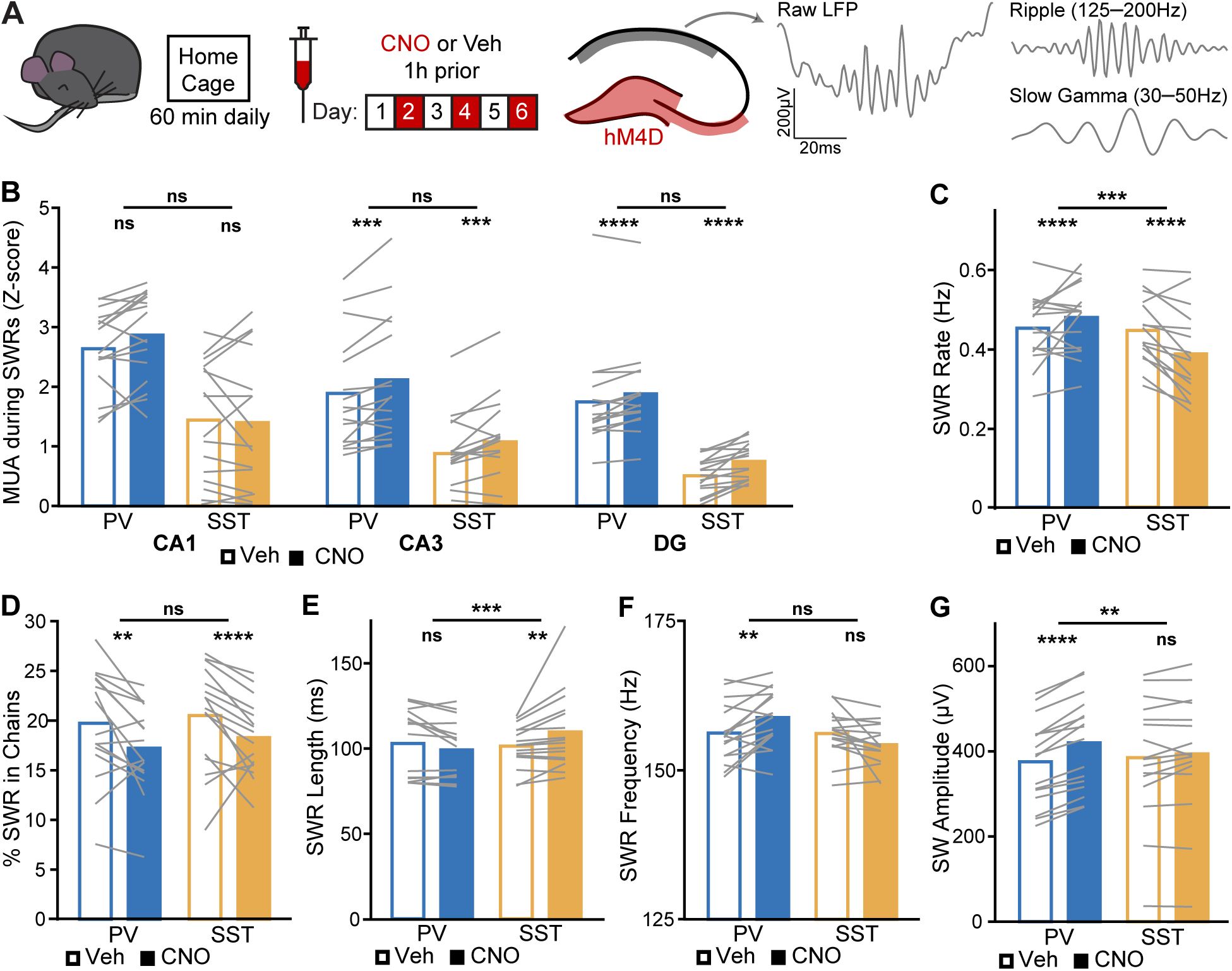
**Suppressing PV^+^ or SST^+^ interneurons bidirectionally modulates SWR signatures of CA3 coupling to CA1.** (A) Mice were recorded during sleep and awake rest over 6 daily home cage sessions, alternating vehicle and CNO treatment. Interneurons in DG and CA3 (cyan) were inhibited while SWRs and related oscillations were assessed in CA1 stratume pyramidale (pyr) and stratum radiatum (sr) (magenta). Representative raw, ripple filtered, and SG filtered traces of a SWR event from a CA1- pyr site of a PV-Cre/SST-Cre mouse during vehicle treatment. (B) Normalized recruitment to SWRs of MUA in CA1 (PV: p = 0.13; SST: p = 0.46; PV vs SST: p = 0.023), CA3 (PV: p = 0.00075; SST: p = 0.00076; PV vs SST: p = 0.095), and DG (PV: p = 5.5×10^-6^; SST: p = 3.4×10^-5^; PV vs SST: p = 0.059). (C) SWR rate (PV: p = 1.9×10^-6^; SST: p = 4.6×10^-5^; PV vs SST: p = 0.00098). (D) Percent of SWRs following another SWR within 200 ms (PV: p = 0.001; SST: p = 2.5×10^-5^; PV vs SST: p = 0.99). (E) SWR temporal length (PV: p = 0.033; SST: p = 0.0023; PV vs SST: p = 0.00034). (F) SWR instantaneous frequency (PV: p = 0.0012; SST: p = 0.033; PV vs SST: p = 0.018). (G) SW amplitude (PV: p = 6.4×10^-27^; SST: p = 0.25; PV vs SST: p = 0.005). N = 16 PV-Cre and n = 16 SST-Cre mice. Statistical details in Table S2. F test of the LMM for treatment effects, likelihood ratio test for genotype-treatment interaction effects. **p < 0.01; ***p < 0.001; ****p < 0.0001. Central values are means and individual points are mean per animal. See also Figures S2 and 6 and Tables S1–S6.

We began by measuring the effect of interneuron suppression on SWR recruitment and found that suppressing either class increased DG and CA3, but not CA1, MUA activity during SWRs (Figure 2B). Thus, suppressing these interneurons unidirectionally disinhibited local DG and CA3 cells while leaving CA1 spiking intact. As SWRs depend on CA3 input to CA1, one might predict that SWR events themselves would be more prevalent following suppression of either PV^+^ or SST^+^ interneurons.

This was not the case. Instead, we found evidence for opposing modulation of SWRs by PV^+^ and SST^+^ interneurons in DG and CA3. Suppressing PV^+^ interneurons increased SWR rate while suppressing SST^+^ interneurons decreased SWR rate, and these effects were significantly different between genotypes (Figure 2C). This effect was not due to differences in SWR detection or baseline MUA (Table S1). There was also a decrease in the proportion of SWRs that participated in chains (Figure 2D) in both genotypes, indicating that broadly increasing DG and CA3 output influences chaining of SWRs. Finally, we examined fast ripples (Valero et al., 2017) to determine whether the observed changes in SWR rate were due to epileptic activity, as ablation or silencing of CA1 or subiculum interneurons can lead to seizures (Drexel et al., 2017; Spampanato and Dudek, 2017). While we observed no behavioral seizures, we did detect a modest increase in the incidence of rare fast ripples when PV^+^ interneurons were suppressed (Table S1), indicating that increased CA3 drive might increase potentially pathological SWRs.

We then assessed the structure of these SWRs and found further evidence for interneuron subtype-specific effects. SST^+^ interneuron suppression increased SWR length, an effect not seen following PV+ interneuron suppression (Figure 2E). Since EC activity can increase prior to SWRs longer than 100 ms (Oliva et al., 2018), we also looked specifically at the proportion of SWRs longer than 100 ms and found an increase when SST^+^ interneurons were suppressed (% of SWRs > 100 ms: vehicle 38.5% vs CNO 49.6%, paired t test, t(15) = 3.67, p = 0.002). Thus, suppressing DG and CA3 SST^+^ interneurons has effects consistent with facilitated EC input to CA1.

Similar differences were seen for other SWR parameters. PV^+^ interneuron suppression alone increased instantaneous frequency of SWRs in CA1 (Figure 2F), but decreased the instantaneous frequency of SWRs in CA3 (Figure S2A), thus the local and downstream effects of these interneurons can be different. During SWRs, PV^+^ interneuron suppression increased sharp-wave (SW) amplitude in CA1, which was specific for this cell type (Figure 2G). However, SWR size was smaller when PV^+^ interneurons were suppressed (Figure S2B) despite no change in CA1 MUA recruitment. Overall, suppressing PV^+^ interneurons increased SWR signatures of CA3 drive to CA1, while suppressing SST^+^ interneurons decreased these signatures, consistent with a role in external drive to DG and CA3 and thereby facilitating external drive to CA1.

### Suppressing PV^+^ or SST^+^ interneurons bidirectionally modulates SWR-coincident SG signatures of CA3 coupling to CA1

Next, we measured SG power coincident with ripples, which is driven by CA3 (Carr et al., 2012; Gillespie et al., 2016; Ramirez-Villegas et al., 2015, 2018). Based on the results above, we again hypothesized that suppressing PV^+^ interneurons would have similar effects to increasing CA3 drive to CA1, while suppressing SST^+^ interneurons would have similar effects to increasing EC drive to CA1.

Following suppression of PV^+^ interneurons, SG power in CA1 was higher across all rest during CNO epochs (Figure 3A) – a signature of increased CA3 drive (Kemere et al., 2013) – and had higher variance in all 3 subregions (Table S2). We therefore used the mean and SD of SG power calculated from vehicle-treated epochs to z-score SG power during all epochs. Suppressing PV^+^ interneurons led to greater SWR-coincident SG power in CA1, CA3, and DG (Figure 3A, 3B, 3D, and S2C). Interestingly, the extent of CA1 SWR-coincident SG power increase correlated across animals with the extent of SW amplitude increase and the extent of SWR frequency increase (Figure S2D–E), suggesting these features reflect similar underlying mechanisms, as has been suggested previously (Oliva et al., 2018; Stark et al., 2014). These effects were specific to PV^+^ interneurons, as suppressing SST^+^ interneurons did not change SG power either at baseline (Table S1) or during SWRs (Figure 3A, 3C, 3D, and S2C). Similarly, only PV^+^ suppression enhanced the increase in SG coherence between CA1 and CA3 or DG during SWRs, (Figure 3E); this was not merely due to increases in activity as CA1 MUA did not increase (Figure 2B). Overall, suppressing PV^+^ interneurons increased SWR activity, consistent with a facilitation of internally generated CA3 spiking patterns that lead to SWRs, while suppressing SST^+^ interneurons decreased SWR activity, consistent with a role in ungating EC inputs to DG and CA3 and thereby changing DG and CA3 firing patterns in a manner that suppressed SWR activity.

**Figure 3.**
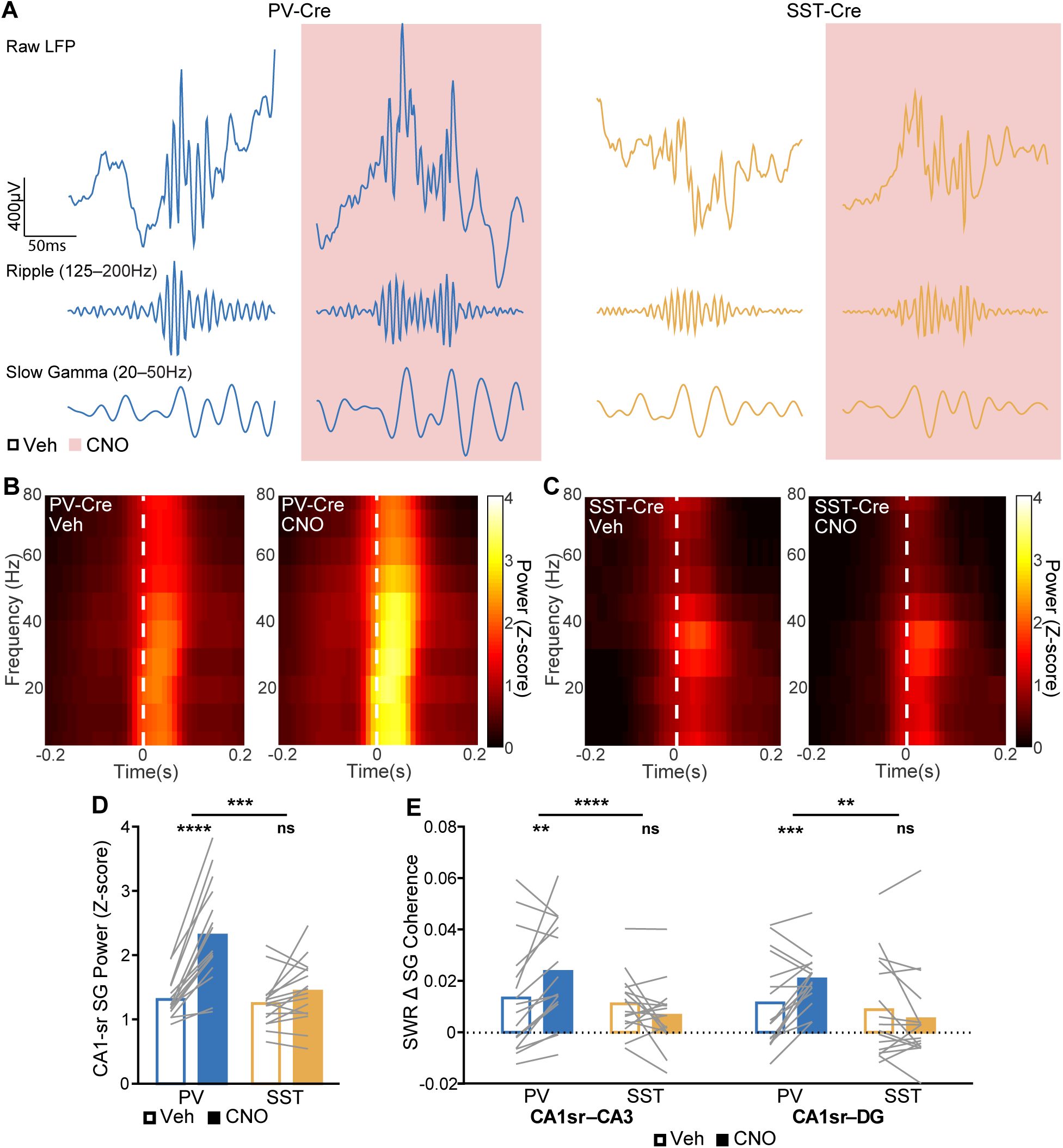
**Suppressing PV^+^ or SST^+^ interneurons bidirectionally modulates SWR-coincident SG signatures of CA3 coupling to CA1.** (A) Example raw (top), SWR-filtered (middle), and SG-filtered (bottom) LFP traces from CA1 pyramidal layer sites from a PV-Cre (left) and an SST-Cre (right) mouse. SG power is higher both outside of and during SWRs in PV-Cre mice following CNO treatment. (B,C) Representative SWR-triggered spectrograms from a CA1-sr layer site during vehicle-treated epochs (left) and CNO-treated epochs (right) in (A) a PV-Cre mouse and (B) an SST-Cre mouse. White dash lines represent threshold crossing for SWR detection. (D) Normalized SG power during SWRs in CA1 (PV: p = 1.7×10^-23^; SST: p = 0.14; PV vs SST: p = 0.00099). (E) Increase during SWRs of SG frequency band coherence between CA1 and CA3 (PV: p = 0.0011; SST: p = 0.17; PV vs SST: p = 9.4×10^-6^) and between CA1 and DG (PV: p = 0.00097; SST: p = 0.43; PV vs SST: p = 0.0036). N = 16 PV-Cre and n = 16 SST-Cre mice. Statistical details in Table S2. F test of the LMM for treatment effects, likelihood ratio test for genotype-treatment interaction effects. **p < 0.01; ***p < 0.001; ****p < 0.0001. Central values are means and individual points are mean per animal. See also Figures S2 and 6 and Tables S1–S6.

### Suppressing PV^+^ or SST^+^ interneurons bidirectionally modulates SG signatures of CA3 coupling to CA1 during movement

We next asked whether these bidirectional effects extended to LFP features observed during movement. We examined CA1 SG, FG, and theta activity during movement (>1 cm/s) in daily 30- minute linear track sessions, focusing on stratum pyramidale (Figure 4A), though similar results were found in recordings from strata radiatum (where CA3 inputs terminate) and lacunosum moleculare (where EC inputs terminate; data not shown). We asked whether suppressing DG and CA3 PV^+^ interneurons ungates internally driven CA3 activity patterns, while suppressing DG and CA3 SST^+^ interneurons ungates EC inputs and thus change firing patterns in a manner that facilitates increasing EC drive to CA1.

**Figure 4.**
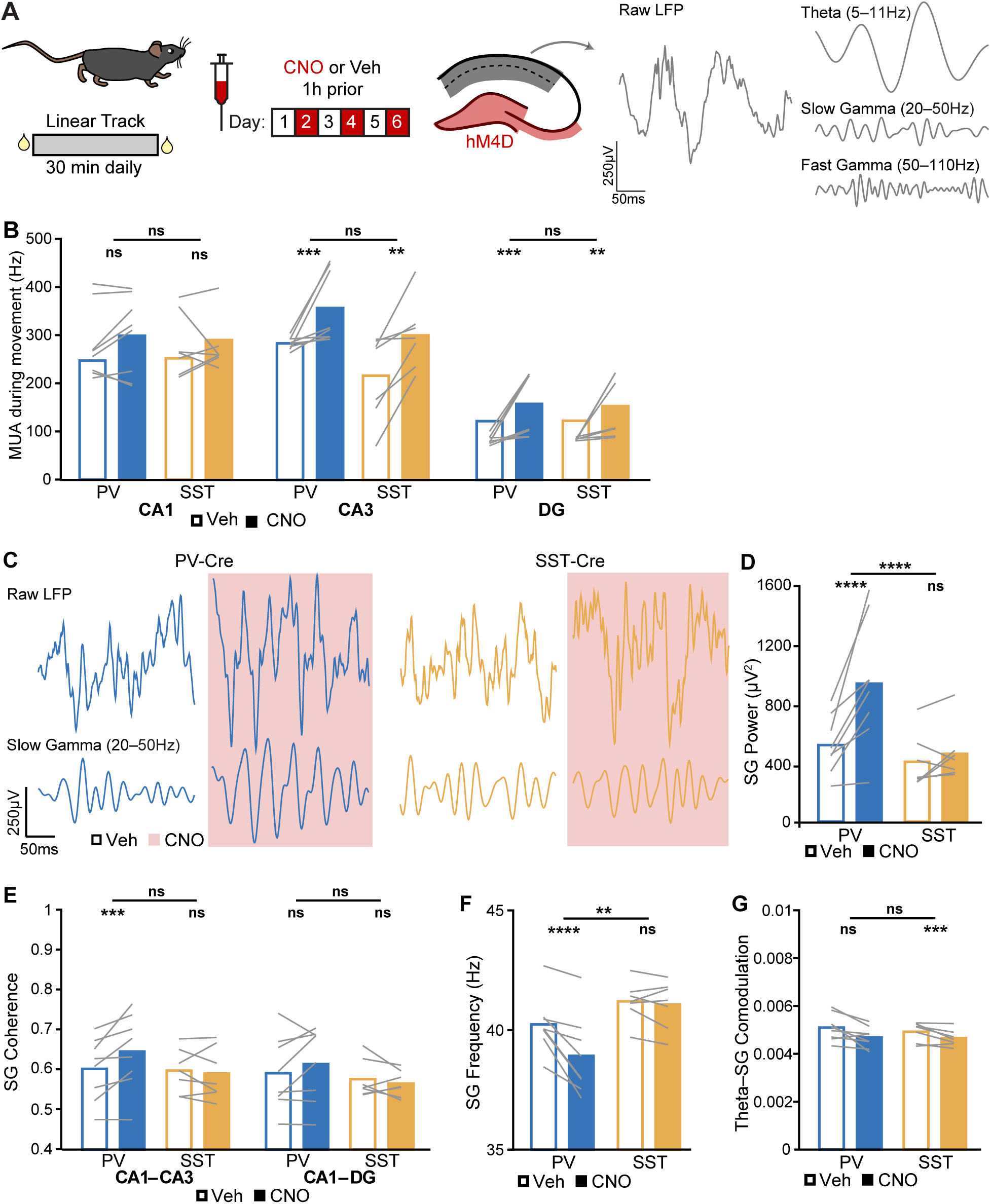
**Suppressing PV^+^ or SST^+^ interneurons bidirectionally modulates SG signatures of CA3 coupling to CA1 during movement.** (A) Mice were recorded during linear track runs over 6 daily sessions, alternating vehicle and CNO treatment. Interneurons in DG and CA3 (cyan) were inhibited while theta and gamma oscillations were assessed in CA1-pyr, CA1-sr, and CA1 stratum lacunosum-moleculare (slm) (magenta). Representative raw trace and LFP filtered for theta, SG, and FG from a CA1-pyr site of a PV-Cre mouse during movement during vehicle treatment. (B) MUA during movement in CA1 (PV: p = 0.72; SST: p = 0.9; PV vs SST: p = 0.34), CA3 (PV: p = 0.00051; SST: p = 0.005; PV vs SST: p = 0.87), and DG (PV: p = 0.00022; SST: p = 0.0063; PV vs SST: p = 0.59). (C) Example raw (top) and SG-filtered (bottom) LFP traces from CA1 pyramidal layer sites from a PV-Cre (left) and an SST-Cre (right) mouse. (D) SG power (PV: p = 2.6×10^-6^; SST: p = 0.13; PV vs SST: p = 4.7×10^-6^). (E) SG frequency band coherence between CA1 and CA3 (PV: p = 0.00028; SST: p = 0.6; PV vs SST: p = 0.56) and between CA1 and DG (PV: p = 0.085; SST: p = 0.3; PV vs SST: p = 0.26). (F) SG instantaneous frequency (PV: p = 2.5×10^-6^; SST: p = 0.56; PV vs SST: p = 0.0061). (G) Theta modulation of SG power (PV: p = 0.12; SST: p = 0.00082; PV vs SST: p = 0.95). N = 8 PV-Cre and n = 7 SST-Cre mice. Statistical details in Table S2. F test of the LMM for treatment effects, likelihood ratio test for genotype-treatment interaction effects. **p < 0.01; ***p < 0.001; ****p < 0.0001. Central values are means and individual points are mean per animal. See also Figures S3, S4, and 6 and Tables S2–S6.

We again began by measuring the effect of interneuron suppression on spiking output and found that suppressing either class increased MUA during movement in DG and CA3, but not CA1 (Figure 4B). Thus, just as for SWR-related activity, suppressing interneurons unidirectionally disinhibited local DG and CA3 cells while leaving CA1 spiking rates intact.

Despite this, we found that PV^+^ and SST^+^ interneurons differentially modulated SG during movement. Suppressing PV^+^ interneurons in DG and CA3 amplified SG power in CA1, but there was no detectable effect following suppression of SST^+^ interneurons. By contrast, suppressing PV^+^ interneurons had no detectable effect on SG power in DG and CA3, while suppressing SST^+^ interneurons dampened SG power locally in DG and CA3. All these effects were significantly different across genotypes (Figure 4B–C and S3A). Relatedly, suppressing PV^+^ interneurons enhanced SG coherence between CA1 and CA3 (Figure 4D). Thus, like during rest periods, suppressing PV^+^ interneurons increased the amplitude of SG in CA1 and the coherence of SG, across regions, consistent with an increase of CA3 drive to CA1. In addition, suppressing PV^+^ interneurons, but not SST^+^ interneurons, reduced SG frequency throughout the hippocampus (Figure 4E and S3B). Finally, suppressing PV^+^ interneurons increased theta modulation of SG only locally in DG and CA3, while suppressing SST^+^ interneurons reduced theta modulation of SG in CA1, and these effects were again detectably different between genotypes. Overall, suppressing PV^+^ interneurons amplified SG while suppressing SST^+^ interneurons had the opposite effect.

### Suppressing PV^+^ or SST^+^ interneurons bidirectionally modulates FG signatures of EC coupling to CA1 during movement

The results above indicate that suppressing PV^+^ or SST^+^ interneurons in DG and CA3 can lead to quite different effects in CA1, despite the increase in DG and CA3 multiunit activity seen in both cases. As described above, a potential for these differences is that these two interneuron classes differentially effect the organization of activity transmitted from CA3 to CA1, and thereby have different effects on CA1. If so, then suppressing SST^+^ interneurons in DG and CA3 would also enhance FG in CA1, despite this typically being characterized as a signature of direct EC input to CA1 (Colgin, 2016). Further, if this hypothesis is correct, then suppressing PV^+^ interneurons in DG and CA3 would have the opposite or no effect, as this would facilitate internally, rather than externally, driven patterns in DG and CA3.

Our data provided strong support for that possibility. Suppressing SST^+^ interneurons amplified FG power in CA1 and DG, an effect specific to this interneuron class (Figure 5A–B and S3D). Interestingly, locally in DG and CA3, suppression of SG in SST-Cre mice appeared to mirror enhancement of FG, where animals with the greatest SG suppression showed the greatest FG enhancement and vice versa (Figure S3E). This suggests that these two signatures of input drive may be opposing, with modulations that decrease SG power leading to greater FG power. Overall, these bidirectional changes in SG and FG power led to decreased SG/FG power ratios in CA1 following SST^+^ suppression and increased SG/FG power ratios throughout the hippocampus following PV^+^ suppression. These SG/FG ratio effects were significantly different between genotypes across the hippocampus (Figure 5C–E and S3F). FG coherence, however, did not change (Figure 5F). In addition, suppressing SST^+^ interneurons increased FG frequency throughout the hippocampus (Figure 5G and S3G). Finally, suppressing SST^+^ interneurons increased theta modulation of FG in CA1 (Figure 5H–J and S3H), an effect which was specific to SST^+^ interneurons. Overall, suppressing SST^+^ interneurons increased CA1 FG activity – a signature of EC drive – while suppressing PV^+^ interneurons had opposing effects. These effects suggest a role for the CA3 input as a modulator of FG in CA1.

**Figure 5.**
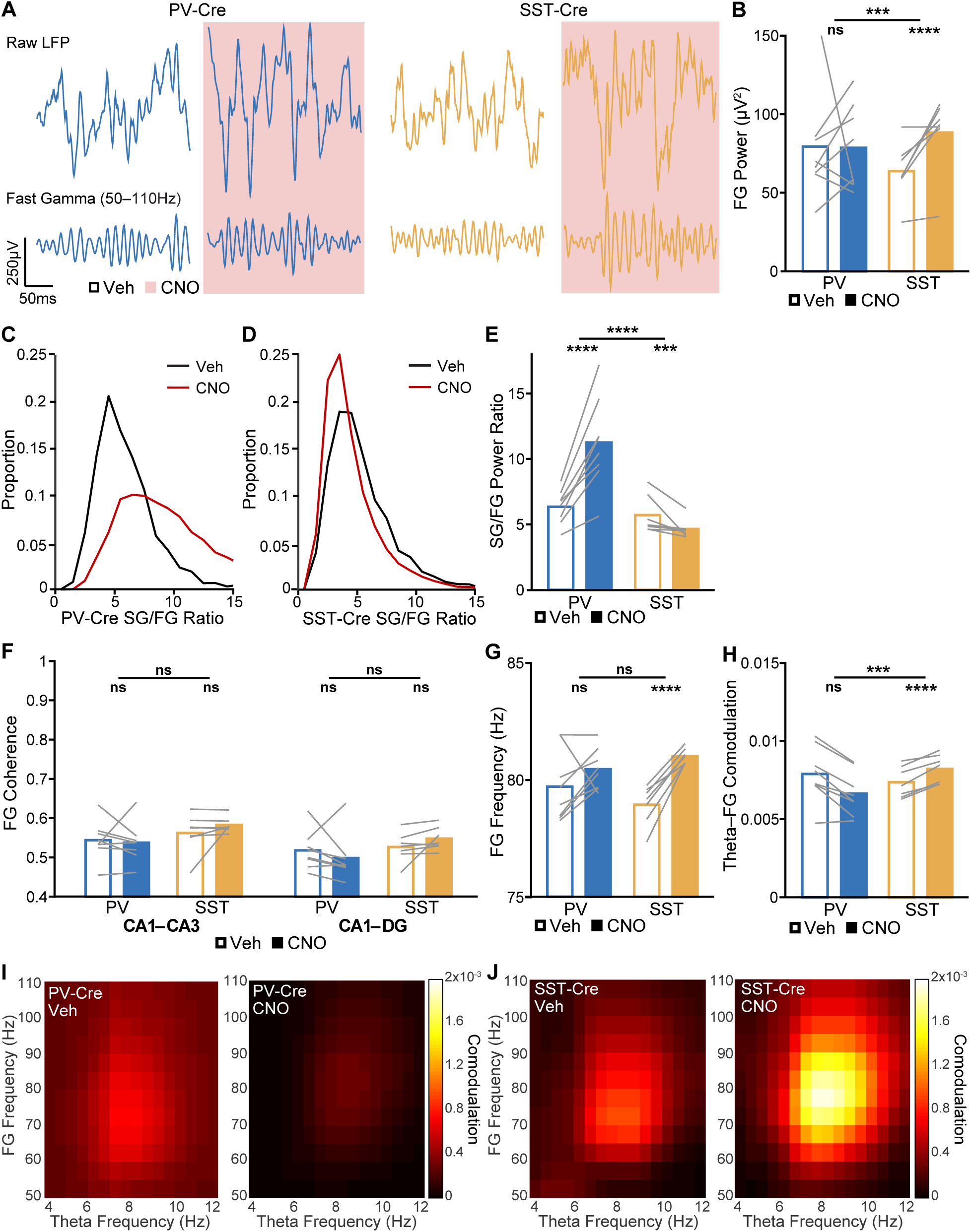
**Suppressing PV^+^ or SST^+^ interneurons bidirectionally modulates FG signatures of EC coupling to CA1 during movement.** (A) Example raw (top) and FG-filtered (bottom) LFP traces from CA1 pyramidal layer sites from a PV-Cre (left) and an SST-Cre (right) mouse. (B) FG power (PV: p = 0.27; SST: p = 3.3×10^-5^; PV vs SST: p = 0.00054). (C,D) Representative distributions of SG/FG power ratios over 1 s bins in (C) a PV-Cre mouse and (D) an SST-Cre mouse. (E) Ratio of SG power to FG power (PV: p = 9.7×10^-25^; SST: p = 0.00013; PV vs SST: p = 7.4×10^-6^). (F) FG frequency band coherence between CA1 and CA3 (PV: p = 0.83; SST: p = 0.2; PV vs SST: p = 0.24) and between CA1 and DG (PV: p = 0.27; SST: p = 0.076; PV vs SST: p = 0.47). (G) FG instantaneous frequency (PV: p = 0.06; SST: p = 4.3×10^-17^; PV vs SST: p = 0.065). (H) Theta modulation of FG power (PV: p = 0.0098; SST: p = 8×10^-20^; PV vs SST: p = 0.0001). (I,J) Representative comodulograms during vehicle-treated epochs (left) and CNO-treated epochs (right) in a (I) a PV-Cre mouse and (J) an SST-Cre mouse. N = 8 PV-Cre and n = 7 SST-Cre mice. Statistical details in Table S2. F test of the LMM for treatment effects, likelihood ratio test for genotype-treatment interaction effects. **p < 0.01; ***p < 0.001; ****p < 0.0001. Central values are means and individual points are mean per animal. See also Figures S3, S4, and 6 and Tables S2–S6.

### Suppressing PV^+^ or SST^+^ interneurons unidirectionally modulates theta during movement

Next, we asked whether PV^+^ and SST ^+^ interneurons might regulate theta. Previous work has suggested that theta throughout the hippocampus is driven by EC and MS inputs, but not by CA3 inputs (Colgin, 2016; Middleton and McHugh, 2016). Consequently, we predicted that suppressing SST^+^ interneurons would enhance theta power in DG and CA3 only, perhaps through ungating EC inputs, while suppressing PV^+^ interneurons would have no effect. Contrary to our hypothesis, suppressing either PV^+^ or SST^+^ interneurons increased theta power in CA1 but not elsewhere (Figure S4A–B). Neither interneuron type modified theta coherence (Figure S4C) or theta frequency (Figure S4D) save for an increase in theta frequency in PV-Cre mice in DG. These findings suggest that a general increase in CA3 spiking output increases CA1 theta power.

Overall, suppressing PV^+^ interneurons in DG and CA3 increased measures of CA3 coupling onto CA1, including enhancing SG power, coherence, and theta comodulation and dampening FG-theta comodulation. In contrast, suppressing SST^+^ interneurons in DG and CA3 increased measures of EC coupling onto CA1, including enhancing FG power and theta coupling and dampening SG power and theta coupling. Suppressing either interneuron type enhanced CA1 theta power.

### Suppressing both PV^+^ and SST^+^ interneurons increases CA3 coupling onto CA1

Lastly, we assessed the effect of suppressing both interneuron populations simultaneously in PV-Cre/SST-Cre mice (Figure S5 and S6 and Table S4). Ablating AMPA currents to all hippocampal interneurons increases SWR rate and frequency (Caputi et al., 2012), similar to DG and CA3 PV^+^ interneuron suppression (Figure 2C and 2F). Thus, we hypothesized that suppressing both interneuron types would follow suppressing only PV^+^ interneurons. In all measures except for comodulation of SG by theta in DG and CA3, we were unable to detect a difference between suppressing both interneuron types and suppressing PV+ interneurons only. In this instance, suppressing both interneuron types decreased theta modulation of SG, an effect not observed when suppressing either type alone. In contrast, we were able to detect several differences between suppressing both interneuron types and suppressing SST+ interneurons only. FG frequency and CA1 FG power are notable exceptions, in which case suppressing both interneuron types followed the direction of suppressing SST+ interneurons alone.

### Consistency of effect across genotypes and sexes

To assess the robustness of these effects, we first assessed both males and females, since females have more DG interneurons than males at this age (Leung et al., 2012). Females had slightly higher SWR sizes than males during vehicle treatment epochs (Table S5), but there were otherwise no sex differences. The effect of suppressing interneurons was in the same direction and magnitude in both sexes on most features (Table S6) with a few exceptions. Suppressing SST^+^ interneurons decreased SW amplitude in females but had the opposite effect in males, leading to no effect overall, and enhanced FG power to a greater degree in females than in males. Interestingly, suppressing either or both interneuron classes led to a greater increase in DG and CA3 MUA during movement in females; though it did not have functional consequences, this underlines the need for sex-specific analyses in manipulation experiments. With those minor exceptions, the observed effects do not appear to be driven mainly in animals of a particular sex.

We further verified that differences in LFP during movement were not due to differences in running speed (vehicle 4.8 cm/s vs CNO 5.1 cm/s, paired t test, t(29) = 1.31, p = 0.2). Then, we confirmed that none of these effects could be confounded by differences between genotypes during vehicle treatment sessions (Table S3). We did note, however, that PV-Cre had different DG and CA3 SG frequencies and SG/FG ratios from SST-Cre mice during vehicle treatment (Table S3), but since the direction of this difference was opposite from the direction of treatment effect, it does not affect our interpretation. Finally, to control for possible off-target effects of CNO, we treated mice injected with an empty vector as a control and observed no differences between vehicle and CNO treated epochs (Figure S5 and S6 and Table S4).

## DISCUSSION

We carried out targeted inhibition of PV^+^ or SST^+^ interneurons in the DG and CA3 regions of the hippocampus and found clear evidence for differential effects on activity in downstream CA1. Our findings provide evidence that PV^+^ and SST^+^ interneurons in DG and CA3 differentially regulate information flow through the hippocampus: suppression of PV^+^ interneurons increased signatures of internal drive associated with CA3 input to CA1 while suppression of SST^+^ interneurons increased signatures of external drive associated with EC input to CA1 (Figure 6). These bidirectional effects were observed despite both interneuron classes increasing the spiking output of DG and CA3 without changing spiking rates in CA1.

**Figure 6.**
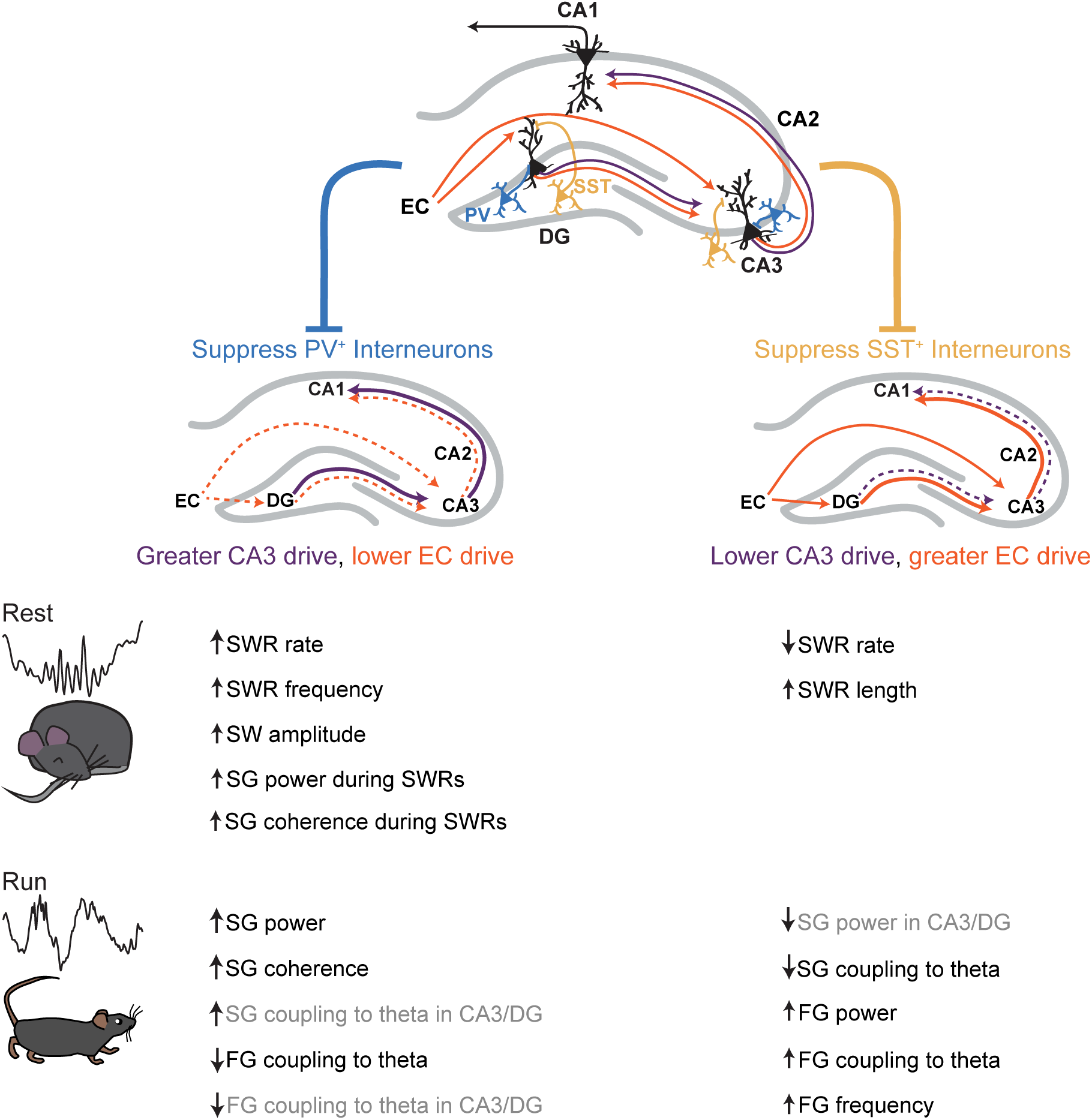
**PV^+^ and SST^+^ interneurons in DG and CA3 bidirectionally modulate signatures of internal and external drive.** CA1 receives direct input from CA3 and indirect input from DG via CA3, and these regions in turn receive projections from EC layer II. PV^+^ (blue) and SST^+^ (yellow) interneurons in DG and CA3 appear to regulate the switch between greater CA3 drive to CA1 and greater EC drive to CA1. When PV^+^ interneurons are suppressed, signatures of CA3 drive are enhanced and signatures of EC drive are curtailed. When SST^+^ interneurons are suppressed, signatures of CA3 drive are curtailed and signatures of EC drive are enhanced. These manifest as changes to SWR, SW, and SG properties during rest periods and to theta, SG, and FG during running periods both downstream in CA1 (black) or locally in DG and CA3 (grey). These interneurons may contribute to the modulation between these internal and external inputs to modulate memory representations.

Specifically, suppressing PV^+^ interneurons led to SWR and SG/FG modulation consistent with increasing CA3 drive and decreasing EC drive to CA1: SWRs seen during immobility had higher incidence, faster frequency, greater amplitude of coincident SWs and SG, and showed higher SG coherence across regions; SG during movement had greater power and coherence across regions while FG during movement had reduced modulation by theta. These findings concur with two recent studies which modulated CA3 PV^+^ interneurons in rats and *ex vivo* (Antonoudiou et al., 2020; López-Madrona et al., 2020). By contrast, suppressing SST^+^ interneurons led to SWR and SG modulation consistent with decreasing CA3 drive and increasing EC drive to CA1: SWRs had lower incidence and greater length, while SG during movement had reduced modulation by theta and FG during movement had greater power. Changes in LFP signatures in DG and CA3 were largely consistent with CA1, suggesting these patterns of activity changed locally and then propagated.

These roles are remarkably similar to those identified for PV^+^ and SST^+^ interneurons in CA1, where PV^+^ interneurons regulate CA3 drive and SST^+^ interneurons regulate EC drive (Fernández-Ruiz et al., 2017; Udakis et al., 2020). We therefore hypothesize that across the hippocampus, PV^+^ and SST^+^ interneurons are likely to be modulated coherently as different flows of information are needed. Importantly, clear differences were observed following inhibition of the two interneuron subtypes despite the existence of substantial variation within interneuron classes (Harris et al., 2018; Klausberger and Somogyi, 2008) and the presence of PV and SST co-expressing cells (Jinno and Kosaka, 2000), suggesting overarching roles for PV^+^ and SST^+^ interneurons in DG and CA3 in regulating information flow through the hippocampal network.

Our findings also identify a dissociation between overall activity levels in CA3 and signatures of CA3 drive in CA1. Suppressing SST^+^ interneurons, which we hypothesized would ungate EC inputs to DG and CA3, led to an increase CA3 multiunit activity. Despite that increase, we observed a decrease in signatures of CA3 input to CA1 but an increase in FG, a signature of direct EC input to CA1 (Colgin, 2016). Importantly, this dissociation between activity and influence is are consistent with the observation of reduced FG signatures of EC drive to CA1 following CA3 silencing (Middleton and McHugh, 2016). These findings suggest that FG in CA1, rather than a pure signature of EC drive to CA1, is best understood as a signature of overall EC drive to both CA1 and the upstream DG and CA3 network. More broadly, these findings highlight the likely importance of specific patterns of CA3 spiking in influencing CA1: suppression of both interneuron types lead to increases in CA3 output but opposite effects on CA3-versus EC-associated LFP patterns in CA1. This is most likely a result of the engagement of different patterns of CA3 spiking.

In controlling input drive, these interneuron classes may regulate hippocampal information processing. The hippocampus contributes to encoding, consolidation, and retrieval of memories, and is thought to alternate between three distinct network states in order to do so (Buzsáki, 1989; Kay and Frank, 2019; Sosa et al., 2018). Encoding of new information is driven by external inputs from the EC, consolidation of previous information by internal inputs from the CA3, and retrieval by a combination of EC and CA3 inputs (Carr and Frank, 2012). Thus, inteneuron-mediated alterations in the strengths of these two inputs could help support different memory functions. Consistent with this possibility, previous behavioral studies examining DG and CA3 PV^+^ and SST^+^ interneurons during contextual threat conditioning support the hypothesis that PV^+^ interneuron suppression could support consolidation by ungating internal CA3 input while SST^+^ interneuron suppression could support encoding by ungating external EC input (Guo et al., 2018; Stefanelli et al., 2016; Zou et al., 2016). In sum, the hippocampal circuit may engage or disengage PV^+^ and SST^+^ interneurons during appropriate learning phases to alter the balance between CA3 and EC inputs and thereby support consolidation or encoding processes.

Previous studies of the CA1 LFP signatures further illustrate how DG and CA3 PV^+^ and SST^+^ interneurons might modulate learning and memory processes. First, suppressing SST^+^ interneurons reduced SWR rate, which might reduce learning, as abolishing SWRs disrupts learning (Ego-Stengel and Wilson, 2010; Girardeau et al., 2009; Jadhav et al., 2012) and lower SWR rate predicts memory impairments in Alzheimer’s models(Jones et al., 2019). However, we found that suppressing SST^+^ interneurons also lengthened SWRs, which may increase the length of the underlying replay trajectory and improve working memory (Fernández-Ruiz et al., 2019). In contrast, suppressing PV^+^ interneurons increased fast ripple incidence, which may be a pathological conversion from SWRs (Behrens et al., 2007; Foffani et al., 2007) which in turn could disrupt normal encoding (Ewell et al., 2019). However, suppressing PV^+^ interneurons heightened the extent to which SG coherence increased during SWRs, which is associated with greater replay fidelity (Carr et al., 2012). Finally, suppressing PV^+^ interneurons increased SG power, which could support retrieval following task learning (Muzzio et al., 2009; Tort et al., 2009), while suppressing SST^+^ interneurons increased FG power and modulation by theta, which could support encoding of new locations (Zheng et al., 2016) and working memory maintenance (Axmacher et al., 2010). Future experiments should examine if there is a casual relationship between activity of these GABAergic populations and spatial task performance.

SST^+^ interneurons in the DG are specifically lost in Alzheimer’s disease and normal aging models, and the extent of their loss correlates with memory impairments (Andrews-Zwilling et al., 2010; Leung et al., 2012; Spiegel et al., 2013). These models show reduced SWR rate, frequency, and coincident SG power (Cayzac et al., 2015; Ciupek et al., 2015; Cowen et al., 2018; Gillespie et al., 2016; Iaccarino et al., 2016; Nicole et al., 2016; Wiegand et al., 2016), and the extent of these impairments predicts memory deficits (Jones et al., 2019). Suppression of DG and CA3 SST^+^ interneurons induces similar changes in SWR characteristics, serving as further evidence that loss of these interneurons may be directly responsible for SWR alterations in pathological and normal aging and related memory deficits.

Beyond the hippocampus, many other brain regions facilitate multiple information processing roles through a balance of inputs. This could also be modulated by PV^+^ and SST^+^ interneurons, as they have been shown to have distinct and often bidirectional roles in cortex. Specifically, this is found in tuning (Miao et al., 2017; Wilson et al., 2012), beta and theta frequency activity (Chen et al., 2017), stimulus-induced gamma rhythms (Hakim et al., 2018; Veit et al., 2017), slow waves and spindles (Funk et al., 2017; Kuki et al., 2015; Niethard et al., 2018), NREM (Funk et al., 2017), reward encoding (Kvitsiani et al., 2013), and working memory (Abbas et al., 2018; Kim et al., 2016). These results are broadly consistent with our results showing bidirectional modulation of signatures of internal and external inputs by DG and CA3 PV^+^ and SST^+^ interneuron populations, and we suggest that the distinction between internal and external drive may be useful in understanding the roles of these interneurons outside the hippocampus.

## ACKNOWLEDGEMENTS

This work was supported by the National Science Foundation Graduate Research Fellowship No. 1144247, National Institute on Aging Predoctoral Fellowship No. F31AG057150, and Genentech Foundation Fellowship to E.A.A.J, funding from the Howard Hughes Medical Institutes to L.M.F., and National Institute on Aging grants RF1AG047655, RF1AG055421, and R01AG055682 to Y.H. We thank Eric Denovellis for assistance with designing the linear mixed effects model, and Max Liu and Scott Owen for assistance with linear track construction, and Alex Gonzalez and Abhilasha Joshi for manuscript feedback.

## AUTHOR CONTRIBUTIONS

E.A.A.J., Y.H., and L.M.F. designed and coordinated the study. E.A.A.J. carried out most studies, performed all data analysis, and created all figures. E.A.A.J. and L.M.F wrote the manuscript. A.K.G. and A.R. assisted with *in vivo* electrophysiological recordings. M.Z. and B.D. performed *ex vivo* electrophysiology experiments. A.R., N.K., M.N., and H.Y. assisted with immunohistochemistry. S.Y.Y. managed mouse lines and performed perfusions. K.H. and T.M.G. assisted with behavioral task design and provided advice on data analysis. Y.H. and L.M.F. provided advice on data analysis and interpretations, edited the manuscript, and supervised the project.

## DECLARATION OF INTERESTS

Y.H. is a co-founder and scientific advisory board member of E-Scape Bio, Inc., GABAeron, Inc., and Mederon Bio, LLC. Other authors declare no competing financial interests.

---

**Figure S1.**
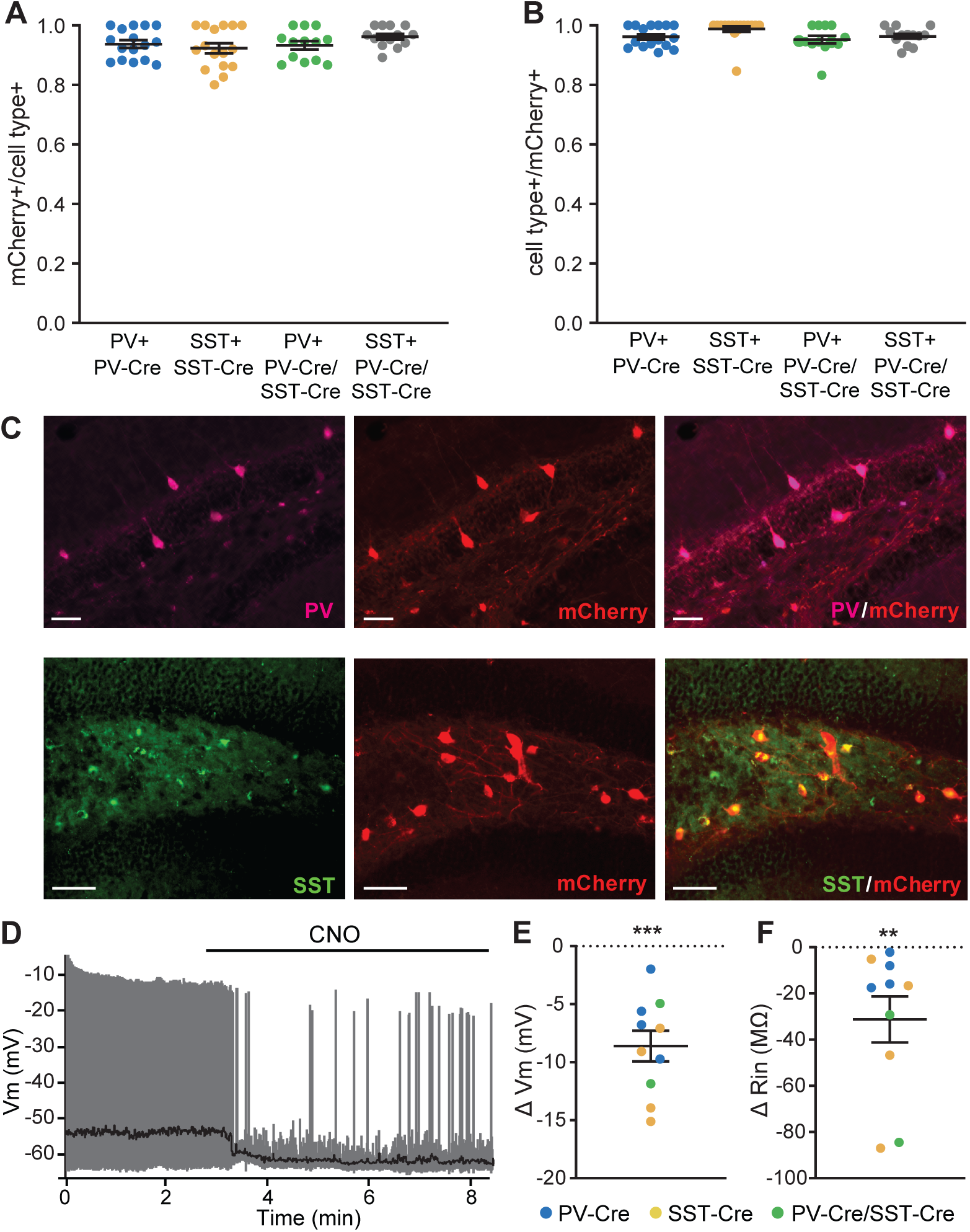
**DREADDs function and expression in PV^+^ and SST^+^ interneurons in DG and CA3. Related to Figure 1.** (A, B) Proportion of (A) mCherry^+^ cells in PV^+^ and SST^+^ cells and (B) PV^+^ and SST^+^ cells in mCherry^+^ cells. N = 16 PV-Cre mice, n = 16 SST-Cre mice, and n = 13 PV-Cre/SST-Cre mice. (C) Example of mCherry coexpression with PV (top) and SST (bottom) in DG. Scale bars are 50 μm. (D) CNO application to *ex vivo* hippocampal sections from a PV-Cre animal reduces the firing rate and resting membrane potential (black trace) of a PV^+^ interneuron expressing hM4D. (E,F) CNO application to *ex vivo* hippocampal section with PV^+^ and SST^+^ interneurons expressing hM4D (D) hyperpolarizes cells (1-sample t test, t(9)=6.57, p = 0.0001) and (E) increases input resistance (1-sample Wilcoxon test, W = -55, p = 0.002). N = 4 PV-Cre slices, n = 4 SST-Cre slices, and n = 2 PV-Cre/SST-Cre slices. Error bars are mean ± SEM. ***p < 0.001.

**Figure S2.**
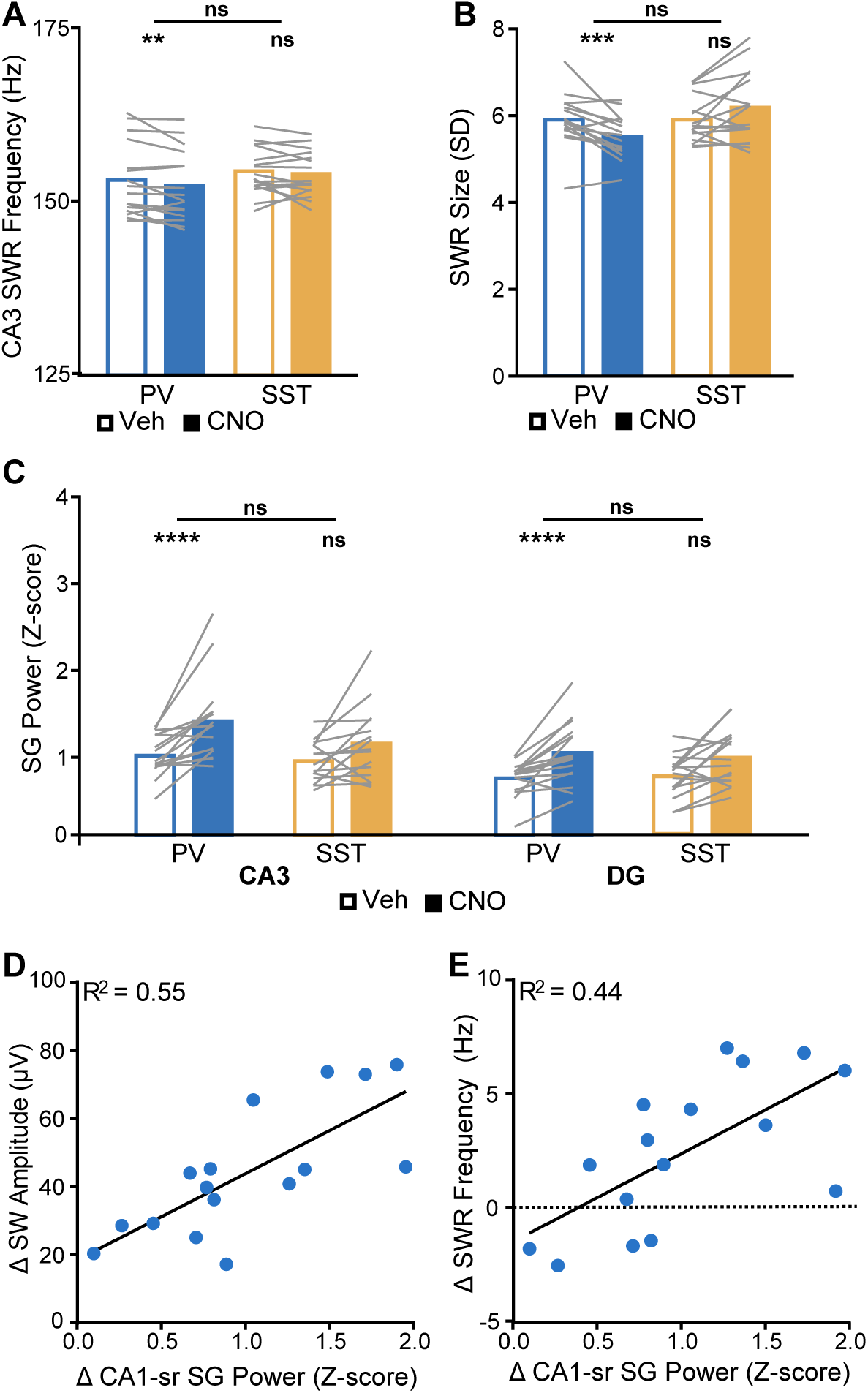
**Additional properties of SWRs during interneuron suppression. Related to Figures 2 and 3.** (A) SWR instantaneous frequency in CA3 (PV: p = 0.0032; SST: p = 0.45; PV vs SST: p = 0.47). (B) SWR size (PV: p = 0.00048; SST: p = 0.07; PV vs SST: p = 0.011). (C) Normalized SG power during SWRs in CA3 (PV: p = 2×10^-5^; SST: p = 0.068; PV vs SST: p = 0.96) and DG (PV: p = 2.2×10^-5^; SST: p = 0.0058; PV vs SST: p = 0.95). (D,E) Change SWR-coincident SG power in CA1 in PV-Cre animals upon CNO treatment predicts (D) change in SW amplitude (F(1,14) = 17.29, p = 0.001) and (E) change in SWR frequency (F(1,14) = 10.97, p = 0.0051). Pearson correlation. N = 16 PV-Cre and n = 16 SST-Cre mice. Statistical details in Table S2. F test of the LMM for treatment effects, likelihood ratio test for genotype-treatment interaction effects. **p < 0.01; ***p < 0.001; ****p < 0.0001. See also Tables S1–S6.

**Figure S3.**
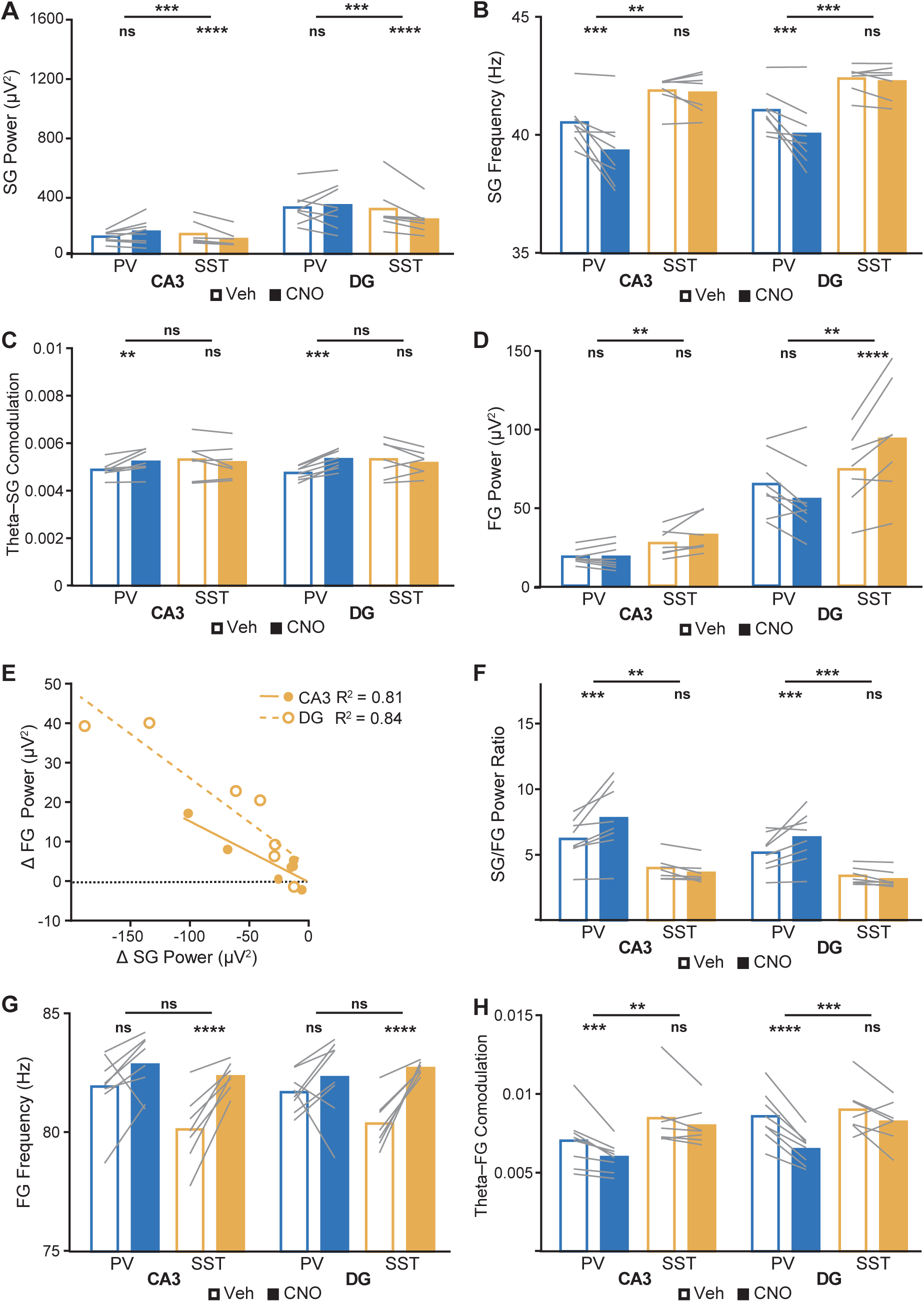
**Properties of SG and FG in DG and CA3 during interneuron suppression. Related to Figures 4 and 5.** (A) SG power in CA3 (PV: p = 0.15; SST: p = 5.2×10^-5^; PV vs SST: p = 0.00088) and DG (PV: p = 0.5; SST: p = 1.3×10^-7^; PV vs SST: p = 0.00042). (B) SG instantaneous frequency in CA3 (PV: p = 0.00023; SST: p = 0.8; PV vs SST: p = 0.0025) and DG (PV: p = 0.0005; SST: p = 0.37; PV vs SST: p = 0.00015). (C) Theta modulation of SG power in CA3 (PV: p = 0.0011; SST: p = 0.16; PV vs SST: p = 0.0049) and DG (PV: p = 0.0003; SST: p = 0.22; PV vs SST: p = 0.0054). (D) FG power in CA3 (PV: p = 0.42; SST: p = 0.0076; PV vs SST: p = 0.0027) and DG (PV: p = 0.81; SST: p = 1.9×10^-5^; PV vs SST: p = 0.0025). (E) Change in SG power in SST-Cre animals upon CNO treatment predicts change in FG power in CA3 (F(1,5) = 20.7, p = 0.0061) and DG (F(1,5) = 26, p = 0.0038). Pearson correlation. (F) Ratio of SG power to FG power in CA3 (PV: p = 0.00036; SST: p = 0.0071; PV vs SST: p = 0.002) and DG (PV: p = 0.00031; SST: p = 0.002; PV vs SST: p = 0.00064). (G) FG instantaneous frequency in CA3 (PV: p = 0.02; SST: p = 2.6×10^-9^; PV vs SST: p = 0.12) and DG (PV: p = 0.23; SST: p = 4.1×10^-11^; PV vs SST: p = 0.062). (H) Theta modulation of FG power in CA3 (PV: p = 0.00041; SST: p = 0.19; PV vs SST: p = 0.0026) and DG (PV: p = 7.5×10^-11^; SST: p = 0.11; PV vs SST: p = 0.00053). N = 8 PV-Cre and n = 7 SST-Cre mice. Statistical details in Table S2. F test of the LMM for treatment effects, likelihood ratio test for genotype-treatment interaction effects. **p < 0.01; ***p < 0.001; ****p < 0.0001. Central values are means and individual points are mean per animal. See also Tables S2–S6.

**Figure S4.**
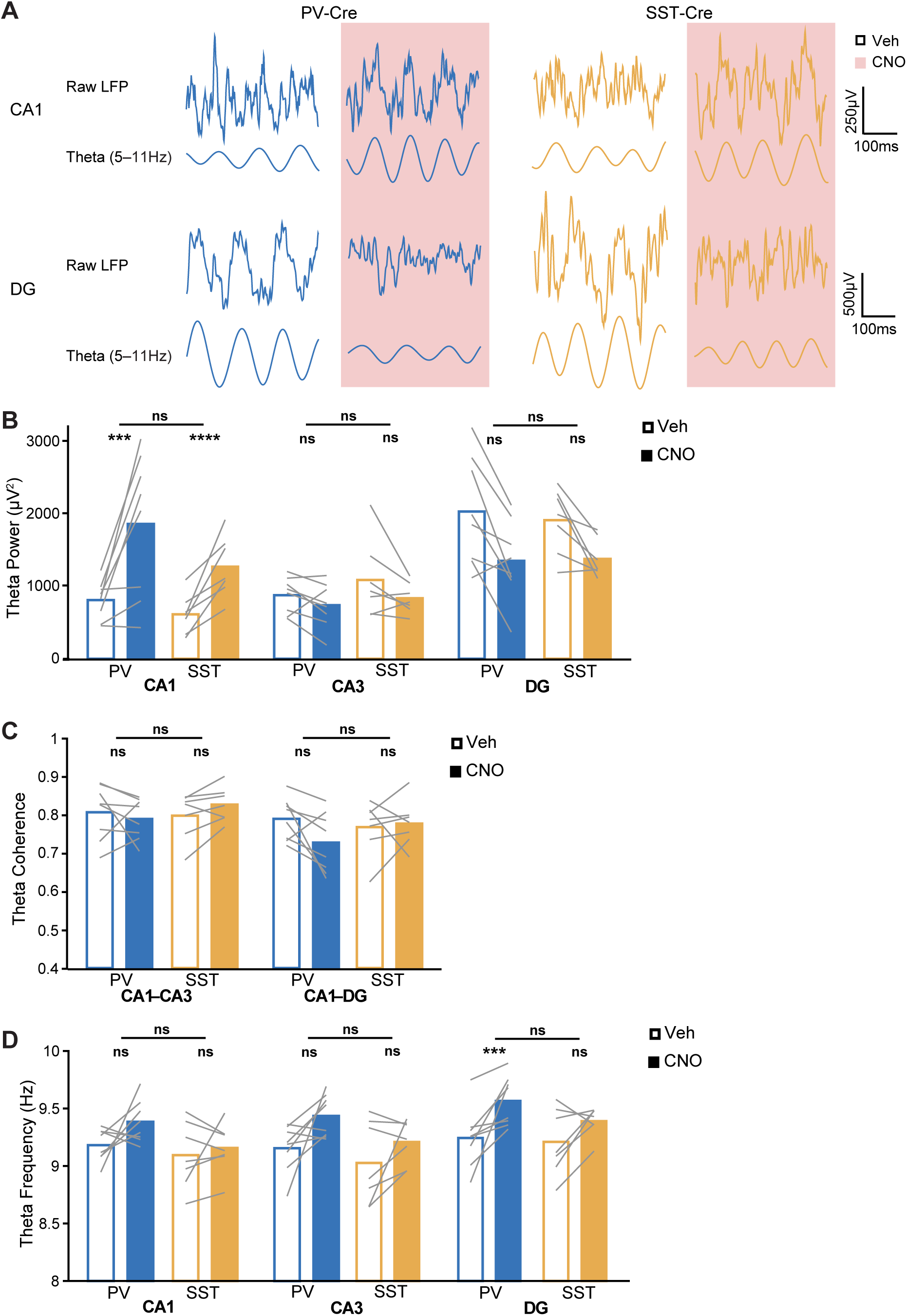
**Suppressing PV^+^ or SST^+^ interneurons unidirectionally modulates theta during movement. Related to Figures 4 and 5.** (A) Example raw and theta-filtered LFP traces from CA1 pyramidal layer (top) and DG granule cell layer (bottom) sites from a PV-Cre (left) and an SST-Cre (right) mouse. (B) Theta power in CA1 (PV: p = 0.00032; SST: p = 4.2×10^-5^; PV vs SST: p = 0.52), CA3 (PV: p = 0.81; SST: p = 0.13; PV vs SST: p = 0.3), and DG (PV: p = 0.12; SST: p = 0.0097; PV vs SST: p = 0.85). (C) Theta frequency band coherence between CA1 and CA3 (PV: p = 0.57; SST: p = 0.047; PV vs SST: p = 0.62) and between CA1 and DG (PV: p = 0.024; SST: p = 0.76; PV vs SST: p = 0.89). (D) Theta instantaneous frequency in CA1 (PV: p = 0.0082; SST: p = 0.29; PV vs SST: p = 0.06), CA3 (PV: p = 0.0077; SST: p = 0.1; PV vs SST: p = 0.3), and DG (PV: p = 0.00061; SST: p = 0.1; PV vs SST: p = 0.67). N = 8 PV-Cre and n = 7 SST-Cre mice. Statistical details in Table S2. F test of the LMM for treatment effects, likelihood ratio test for genotype-treatment interaction effects. **p < 0.01; ***p < 0.001; ****p < 0.0001. Central values are means and individual points are mean per animal. See also Tables S2–S6.

**Figure S5.**
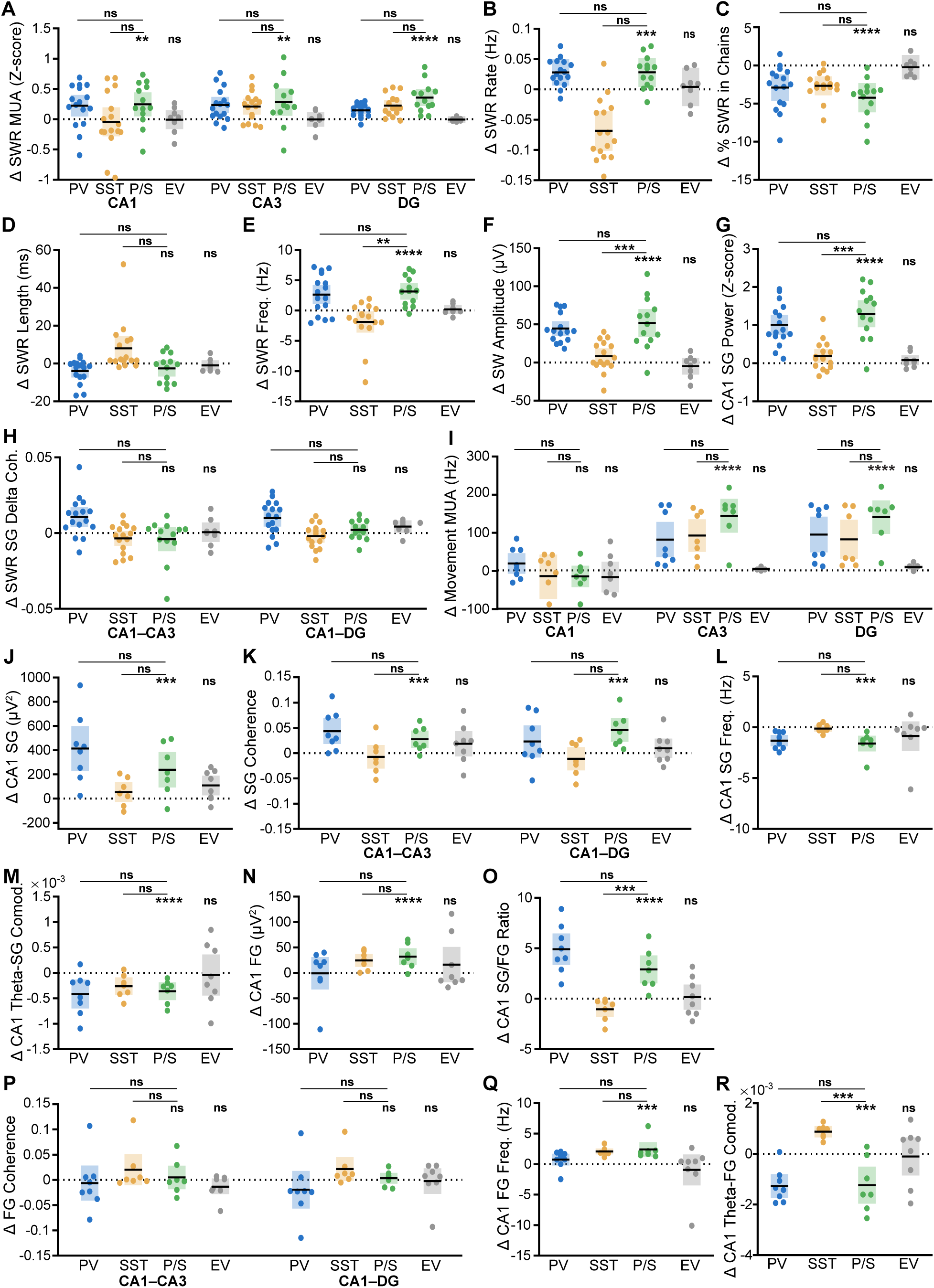
**Suppressing both PV^+^ and SST^+^ interneurons increases CA3 coupling onto CA1. Related to Figures 2–5 and S2–S4.** (A) Normalized recruitment to SWRs of MUA in CA1 (P/S: p = 0.0074; PV vs P/S: p = 1; SST vs P/S: p = 1; EV: p = 0.89), CA3 (P/S: p = 0.0053; PV vs P/S: p = 1; SST vs P/S: p = 1; EV: p = 0.66), and DG (P/S: p = 9×10^-7^; PV vs P/S: p = 1; SST vs P/S: p = 1; EV: p = 0.23). (B) SWR rate (PV-Cre/SST-Cre (P/S): p = 0.00035; PV vs P/S: p = 0.87; SST vs P/S: p = 0.021; Empty Vector (EV): p = 0.74). (C) Percent of SWRs following another SWR within 200 ms (P/S: p = 1.4×10^-5^; PV vs P/S: p = 1; SST vs P/S: p = 0.82; EV: p = 0.77). (D) SWR length (P/S: p = 0.49; PV vs P/S: p = 0.8; SST vs P/S: p = 0.089; EV: p = 0.68). (E) SWR instantaneous frequency (P/S: p = 4.2×10^-6^; PV vs P/S: p = 0.2; SST vs P/S: p = 0.0017; EV: p = 0.62). (F) SW amplitude (P/S: p = 5.2×10^-8^; PV vs P/S: p = 0.066; SST vs P/S: p = 0.00055; EV: p = 0.47). (G) Normalized SG power during SWRs in CA1 (P/S: p = 8.6×10^-12^; PV vs P/S: p = 1; SST vs P/S: p = 0.00089; EV: p = 0.13). (H) Increase during SWRs of SG frequency band coherence between CA1 and CA3 (P/S: p = 0.32; PV vs P/S: p = 1; SST vs P/S: p = 1; EV: p = 0.89 and between CA1 and DG (P/S: p = 0.23; PV vs P/S: p = 1; SST vs P/S: p = 1; EV: p = 0.13). In A–H, n = 16 PV-Cre, n = 16 SST-Cre, n = 13 PV-Cre/SST-Cre and n = 8 empty vector mice. (I) MUA during movement in CA1 (P/S: p = 0.033; PV vs P/S: p = 0.98; SST vs P/S: p = 0.15; EV: p = 0.23), CA3 (P/S: p = 1.1×10^-9^; PV vs P/S: p = 0.51; SST vs P/S: p = 0.89; EV: p = 0.36), and DG (P/S: p = 1.2×10^-9^; PV vs P/S: p = 0.43; SST vs P/S: p = 0.031; EV: p = 0.6). (J) SG power (P/S: p = 0.00065; PV vs P/S: p = 0.78; SST vs P/S: p = 0.73; EV: p = 0.067). (K) SG frequency band coherence between CA1 and CA3 (P/S: p = 0.00058; PV vs P/S: p = 0.6; SST vs P/S: p = 0.45; EV: p = 0.13) and between CA1 and DG (P/S: p = 0.00098; PV vs P/S: p = 0.95; SST vs P/S: p = 0.26; EV: p = 0.3). (L) SG instantaneous frequency (P/S: p = 0.0001; PV vs P/S: p = 0.95; SST vs P/S: p = 0.036; EV: p = 0.28). (M) Theta modulation of SG power (P/S: p = 7.2×10^-7^; PV vs P/S: p = 0.66; SST vs P/S: p = 0.93; EV: p = 0.87). (N) FG power (P/S: p = 1.6×10^-6^; PV vs P/S: p = 0.05; SST vs P/S: p = 1; EV: p = 0.71). (O) Ratio of SG power to FG power (P/S: p = 3.8×10^-6^; PV vs P/S: p = 0.5; SST vs P/S: p = 0.00086; EV: p = 0.92). (P) FG frequency band coherence between CA1 and CA3 (P/S: p = 0.68; PV vs P/S: p = 0.54; SST vs P/S: p = 0.1; EV: p = 0.1) and between CA1 and DG (P/S: p = 0.68; PV vs P/S: p = 0.13; SST vs P/S: p = 0.16; EV: p = 0.82). (Q) FG instantaneous frequency (P/S: p = 0.00012; PV vs P/S: p = 0.51; SST vs P/S: p = 0.0056; EV: p = 0.44). (R) Theta modulation of FG power (P/S: p = 0.00019; PV vs P/S: p = 0.53; SST vs P/S: p = 0.00017; EV: p = 0.91). In I–R, n = 8 PV-Cre, n = 7 SST-Cre, n = 7 PV-Cre/SST-Cre and n = 8 empty vector mice. Statistical details in Table S4. F test of the LMM for treatment effects, likelihood ratio test for genotype-treatment interaction effects. **p < 0.01; ***p < 0.001; ****p < 0.0001. Central values are LMM fixed effect coefficients β ± 95% confidence intervals and individual points are the fitted conditional response for each mouse. See also Figures S5 and Table S4.

**Figure S6.**
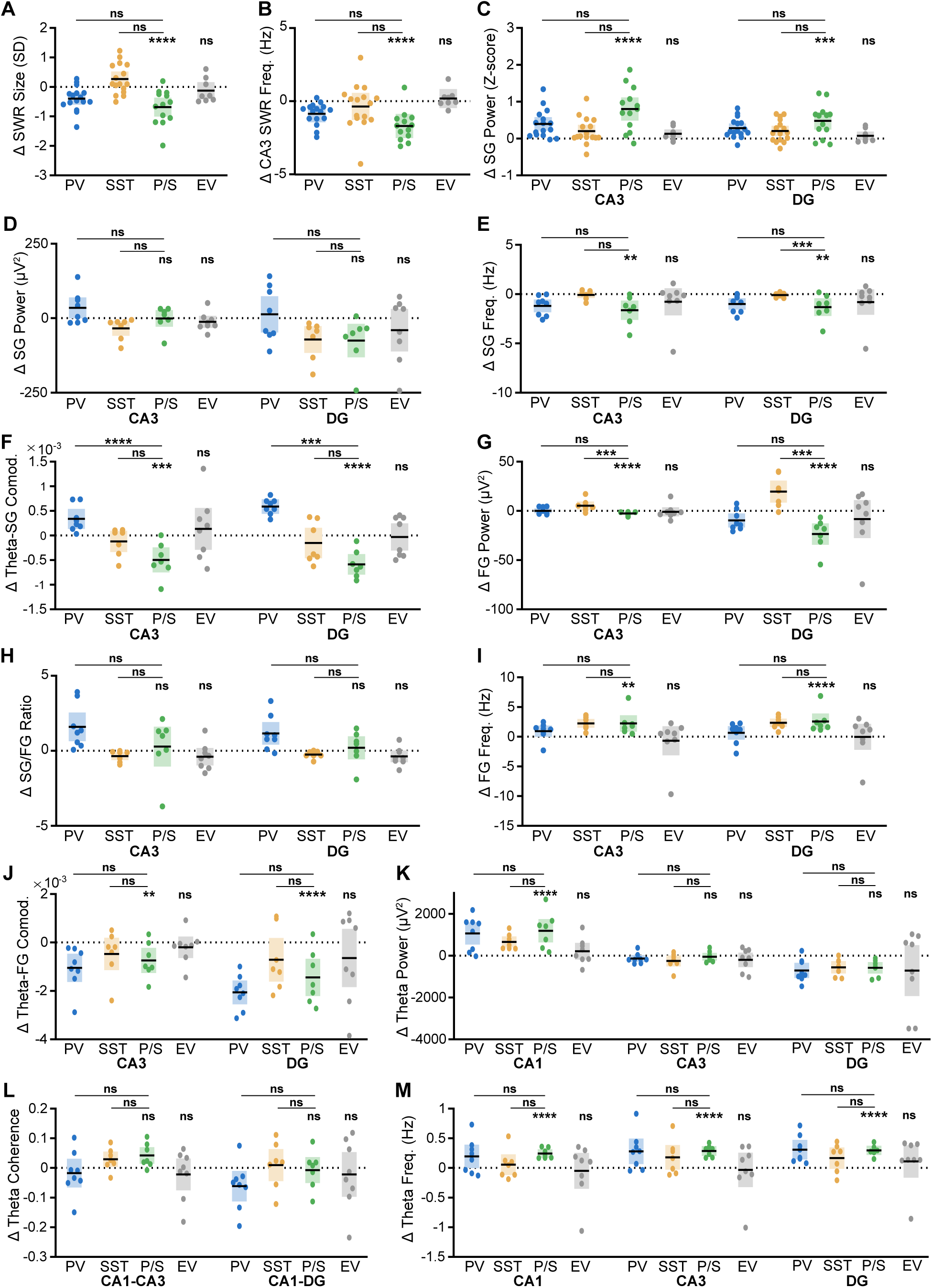
**Additional properties demonstrating suppressing both PV^+^ and SST^+^ interneurons increases CA3 coupling onto CA1. Figures 2–5 and S2–S4.** (A) SWR size (PV-Cre/SST-Cre (P/S): p = 7.2×10^-6^; PV vs P/S: p = 0.56; SST vs P/S: p = 0.031; Empty Vector (EV): p = 0.48). (B) SWR instantaneous frequency in CA3 (P/S: p = 4.3×10^-5^; PV vs P/S: p = 0.9; SST vs P/S: p = 0.45; EV: p = 0.53). (C) Normalized SG power during SWRs in CA3 (P/S: p = 1.1×10^-6^; PV vs P/S: p = 1; SST vs P/S: p = 1; EV: p = 0.062) and DG (P/S: p = 0.00013; PV vs P/S: p = 1; SST vs P/S: p = 1; EV: p = 0.22). In A–C, n = 16 PV-Cre, n = 16 SST-Cre, n = 13 PV-Cre/SST-Cre and n = 8 empty vector mice. (D) SG power in CA3 (P/S: p = 0.84; PV vs P/S: p = 0.47; SST vs P/S: p = 0.49; EV: p = 0.087) and DG (P/S: p = 0.017; PV vs P/S: p = 0.35; SST vs P/S: p = 0.69; EV: p = 0.63). (E) SG instantaneous frequency in CA3 (P/S: p = 0.0029; PV vs P/S: p = 0.7; SST vs P/S: p = 0.012; EV: p = 0.32) and DG (P/S: p = 0.0045; PV vs P/S: p = 0.8; SST vs P/S: p = 0.00055; EV: p = 0.27). (F) Theta modulation of SG power in CA3 (P/S: p = 0.00036; PV vs P/S: p = 7.3×10^-5^; SST vs P/S: p = 0.0048; EV: p = 0.63) and DG (P/S: p = 8.4×10^-11^; PV vs P/S: p = 0.00018; SST vs P/S: p = 0.28; EV: p = 1). (G) FG power in CA3 (P/S: p = 2×10^-6^; PV vs P/S: p = 0.037; SST vs P/S: p = 0.00023; EV: p = 0.36) and DG (P/S: p = 1×10^-10^; PV vs P/S: p = 0.018; SST vs P/S: p = 0.00019; EV: p = 0.64). (H) Ratio of SG power to FG power in CA3 (P/S: p = 0.39; PV vs P/S: p = 0.52; SST vs P/S: p = 0.022; EV: p = 0.23) and DG (P/S: p = 0.29; PV vs P/S: p = 0.66; SST vs P/S: p = 0.012; EV: p = 0.15). (I) FG instantaneous frequency in CA3 (P/S: p = 0.0013; PV vs P/S: p = 0.39; SST vs P/S: p = 0.092; EV: p = 0.51) and DG (P/S: p = 8.6×10^-5^; PV vs P/S: p = 0.061; SST vs P/S: p = 0.0089; EV: p = 0.74). (J) Theta modulation of FG power in CA3 (P/S: p = 0.0024; PV vs P/S: p = 0.68; SST vs P/S: p = 0.79; EV: p = 0.22) and DG (P/S: p = 8.6×10^-5^; PV vs P/S: p = 0.75; SST vs P/S: p = 0.96; EV: p = 0.35). (K) Theta power in CA1 (P/S: p = 2.8×10^-8^; PV vs P/S: p = 0.69; SST vs P/S: p = 0.51; EV: p = 0.1), CA3 (P/S: p = 0.35; PV vs P/S: p = 0.004; SST vs P/S: p = 0.018; EV: p = 0.67), and DG (P/S: p = 0.014; PV vs P/S: p = 0.013; SST vs P/S: p = 0.028; EV: p = 0.61). (L) Theta frequency band coherence between CA1 and CA3 (P/S: p = 0.0064; PV vs P/S: p = 0.66; SST vs P/S: p = 1; EV: p = 0.36) and between CA1 and DG (P/S: p = 0.66; PV vs P/S: p = 0.9; SST vs P/S: p = 0.91; EV: p = 0.59). (M) Theta instantaneous frequency in CA1 (P/S: p = 1.4×10^-8^; PV vs P/S: p = 0.056; SST vs P/S: p = 0.55; EV: p = 0.68), CA3 (P/S: p = 5.7×10^-8^; PV vs P/S: p = 0.16; SST vs P/S: p = 0.86; EV: p = 0.74), and DG (P/S: p = 9.4×10^-12^; PV vs P/S: p = 0.69; SST vs P/S: p = 0.54; EV: p = 0.8). In D–M, n = 8 PV-Cre, n = 7 SST-Cre, n = 7 PV-Cre/SST-Cre and n = 8 empty vector mice. Statistical details in Table S4. F test of the LMM for treatment effects, likelihood ratio test for genotype-treatment interaction effects. **p < 0.01; ***p < 0.001; ****p < 0.0001. Central values are LMM fixed effect coefficients β ± 95% confidence intervals and individual points are the fitted conditional response for each mouse. See also Table S4.

**Table S1.** Effects of suppressing PV^+^ or SST^+^ interneurons on SWR and SG detection. Related to Figures 2, 3, and S2.

| Feature | Layer | Genotype | Veh. Mean | CNO Mean | statistic | df | test value | p |
| --- | --- | --- | --- | --- | --- | --- | --- | --- |
| Ripple Freq. Mean (μV) | CA1-pyr | PV-Cre | 72.39 | 70.96 | Z | NA | 1.6 | 0.1 |
|  |  | SST-Cre | 61.53 | 58.67 | Z | NA | 1.24 | 0.2 |
| Ripple Freq. SD (μV) | CA1-pyr | PV-Cre | 77.3 | 76.69 | t | 15 | 0.32 | 0.8 |
|  |  | SST-Cre | 64.04 | 65.76 | t | 15 | 0.66 | 0.5 |
| Fast Ripple Rate (Hz) | CA1-pyr | **PV-Cre** | **0.0004** | **0.0035** | **Z** | **NA** | **2.93** | **0.003** |
|  |  | SST-Cre | 0.0015 | 0.0013 | Z | NA | 1.17 | 0.2 |
| Rest MUA (Hz) | CA1-pyr | PV-Cre | 10.61 | 10.91 | Z | NA | -0.41 | 0.68 |
|  |  | SST-Cre | 12.28 | 11.49 | Z | NA | -0.16 | 0.88 |
|  | CA3-pyr | PV-Cre | 8.1 | 7.4 | Z | NA | 1.71 | 0.09 |
|  |  | SST-Cre | 6.58 | 5.75 | Z | NA | 1.6 | 0.11 |
|  | DG-gc | PV-Cre | 7.05 | 6.51 | Z | NA | 0.98 | 0.33 |
|  |  | SST-Cre | 5.78 | 5.91 | Z | NA | 0.21 | 0.84 |
| Rest SG Freq. Mean (μV<sup>2</sup>) | CA1-sr | **PV-Cre** | **109.51** | **127.6** | **t** | **15** | **4.03** | **0.001** |
|  |  | SST-Cre | 114.49 | 104.57 | t | 15 | 1.8 | 0.09 |
|  | CA3 | PV-Cre | 152.26 | 157.61 | t | 15 | 1.49 | 0.2 |
|  |  | SST-Cre | 140 | 138.21 | t | 15 | 0.6 | 0.6 |
|  | DG | PV-Cre | 353.58 | 352.37 | t | 15 | 0.15 | 0.9 |
|  |  | SST-Cre | 338 | 338.89 | t | 15 | 0.1 | 0.94 |
| Rest SG Freq. SD (μV<sup>2</sup>) | CA1-sr | **PV-Cre** | **109.06** | **226.98** | **Z** | **NA** | **3.52** | **0.0004** |
|  |  | SST-Cre | 130.07 | 130.39 | t | 15 | 0.1 | 0.96 |
|  | CA3 | **PV-Cre** | **161.1** | **252.93** | **t** | **15** | **3.94** | **0.001** |
|  |  | SST-Cre | 158.63 | 180.56 | Z | NA | 1.4 | 0.16 |
|  | DG | **PV-Cre** | **343.52** | **450.84** | **t** | **15** | **3.47** | **0.003** |
|  |  | SST-Cre | 355.87 | 382.82 | t | 15 | 1.4 | 0.19 |

Paired t tests when the test statistic given is t and Wilcoxon matched pairs signed rank test when the test statistic given is Z. Significant comparisons are bolded. N = 16 PV-Cre and n = 16 SST-Cre mice.

**Table S2.**
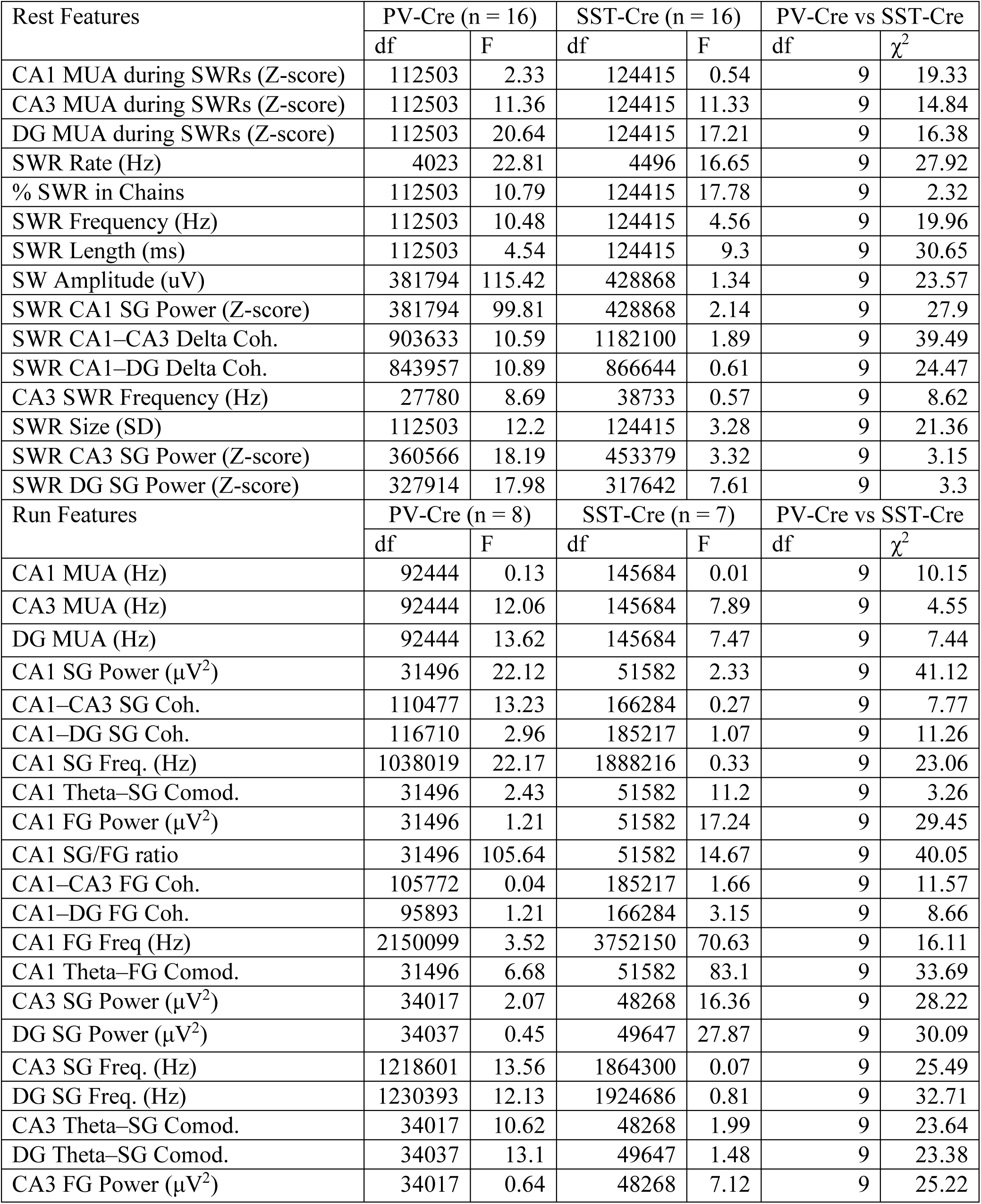
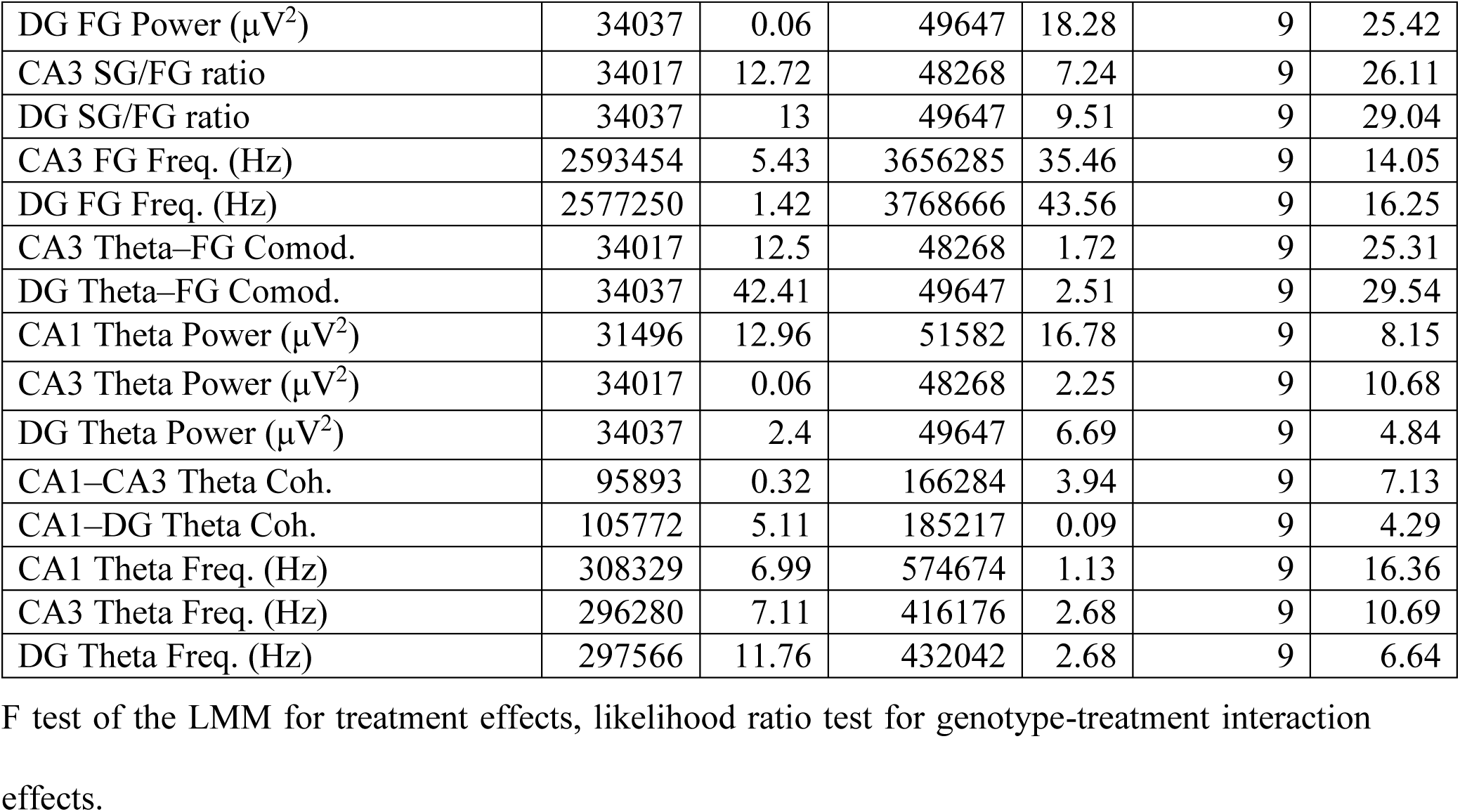
Statistical details for Figures 2–5 and S2–S4.

| Rest Features | PV-Cre (n = 16) |  | SST-Cre (n = 16) |  | PV-Cre vs SST-Cre |  |
| --- | --- | --- | --- | --- | --- | --- |
| | df | F | df | F | df | $\chi^2$ |
| CA1 MUA during SWRs (Z-score) | 112503 | 2.33 | 124415 | 0.54 | 9 | 19.33 |
| CA3 MUA during SWRs (Z-score) | 112503 | 11.36 | 124415 | 11.33 | 9 | 14.84 |
| DG MUA during SWRs (Z-score) | 112503 | 20.64 | 124415 | 17.21 | 9 | 16.38 |
| SWR Rate (Hz) | 4023 | 22.81 | 4496 | 16.65 | 9 | 27.92 |
| % SWR in Chains | 112503 | 10.79 | 124415 | 17.78 | 9 | 2.32 |
| SWR Frequency (Hz) | 112503 | 10.48 | 124415 | 4.56 | 9 | 19.96 |
| SWR Length (ms) | 112503 | 4.54 | 124415 | 9.3 | 9 | 30.65 |
| SW Amplitude (uV) | 381794 | 115.42 | 428868 | 1.34 | 9 | 23.57 |
| SWR CA1 SG Power (Z-score) | 381794 | 99.81 | 428868 | 2.14 | 9 | 27.9 |
| SWR CA1–CA3 Delta Coh. | 903633 | 10.59 | 1182100 | 1.89 | 9 | 39.49 |
| SWR CA1–DG Delta Coh. | 843957 | 10.89 | 866644 | 0.61 | 9 | 24.47 |
| CA3 SWR Frequency (Hz) | 27780 | 8.69 | 38733 | 0.57 | 9 | 8.62 |
| SWR Size (SD) | 112503 | 12.2 | 124415 | 3.28 | 9 | 21.36 |
| SWR CA3 SG Power (Z-score) | 360566 | 18.19 | 453379 | 3.32 | 9 | 3.15 |
| SWR DG SG Power (Z-score) | 327914 | 17.98 | 317642 | 7.61 | 9 | 3.3 |
| Run Features | PV-Cre (n = 8) |  | SST-Cre (n = 7) |  | PV-Cre vs SST-Cre |  |
| | df | F | df | F | df | $\chi^2$ |
| CA1 MUA (Hz) | 92444 | 0.13 | 145684 | 0.01 | 9 | 10.15 |
| CA3 MUA (Hz) | 92444 | 12.06 | 145684 | 7.89 | 9 | 4.55 |
| DG MUA (Hz) | 92444 | 13.62 | 145684 | 7.47 | 9 | 7.44 |
| CA1 SG Power ($\mu V^2$) | 31496 | 22.12 | 51582 | 2.33 | 9 | 41.12 |
| CA1–CA3 SG Coh. | 110477 | 13.23 | 166284 | 0.27 | 9 | 7.77 |
| CA1–DG SG Coh. | 116710 | 2.96 | 185217 | 1.07 | 9 | 11.26 |
| CA1 SG Freq. (Hz) | 1038019 | 22.17 | 1888216 | 0.33 | 9 | 23.06 |
| CA1 Theta–SG Comod. | 31496 | 2.43 | 51582 | 11.2 | 9 | 3.26 |
| CA1 FG Power ($\mu V^2$) | 31496 | 1.21 | 51582 | 17.24 | 9 | 29.45 |
| CA1 SG/FG ratio | 31496 | 105.64 | 51582 | 14.67 | 9 | 40.05 |
| CA1–CA3 FG Coh. | 105772 | 0.04 | 185217 | 1.66 | 9 | 11.57 |
| CA1–DG FG Coh. | 95893 | 1.21 | 166284 | 3.15 | 9 | 8.66 |
| CA1 FG Freq (Hz) | 2150099 | 3.52 | 3752150 | 70.63 | 9 | 16.11 |
| CA1 Theta–FG Comod. | 31496 | 6.68 | 51582 | 83.1 | 9 | 33.69 |
| CA3 SG Power ($\mu V^2$) | 34017 | 2.07 | 48268 | 16.36 | 9 | 28.22 |
| DG SG Power ($\mu V^2$) | 34037 | 0.45 | 49647 | 27.87 | 9 | 30.09 |
| CA3 SG Freq. (Hz) | 1218601 | 13.56 | 1864300 | 0.07 | 9 | 25.49 |
| DG SG Freq. (Hz) | 1230393 | 12.13 | 1924686 | 0.81 | 9 | 32.71 |
| CA3 Theta–SG Comod. | 34017 | 10.62 | 48268 | 1.99 | 9 | 23.64 |
| DG Theta–SG Comod. | 34037 | 13.1 | 49647 | 1.48 | 9 | 23.38 |
| CA3 FG Power ($\mu V^2$) | 34017 | 0.64 | 48268 | 7.12 | 9 | 25.22 |
| DG FG Power ($\mu V^2$) | 34037 | 0.06 | 49647 | 18.28 | 9 | 25.42 |
| CA3 SG/FG ratio | 34017 | 12.72 | 48268 | 7.24 | 9 | 26.11 |
| DG SG/FG ratio | 34037 | 13 | 49647 | 9.51 | 9 | 29.04 |
| CA3 FG Freq. (Hz) | 2593454 | 5.43 | 3656285 | 35.46 | 9 | 14.05 |
| DG FG Freq. (Hz) | 2577250 | 1.42 | 3768666 | 43.56 | 9 | 16.25 |
| CA3 Theta–FG Comod. | 34017 | 12.5 | 48268 | 1.72 | 9 | 25.31 |
| DG Theta–FG Comod. | 34037 | 42.41 | 49647 | 2.51 | 9 | 29.54 |
| CA1 Theta Power ($\mu V^2$) | 31496 | 12.96 | 51582 | 16.78 | 9 | 8.15 |
| CA3 Theta Power ($\mu V^2$) | 34017 | 0.06 | 48268 | 2.25 | 9 | 10.68 |
| DG Theta Power ($\mu V^2$) | 34037 | 2.4 | 49647 | 6.69 | 9 | 4.84 |
| CA1–CA3 Theta Coh. | 95893 | 0.32 | 166284 | 3.94 | 9 | 7.13 |
| CA1–DG Theta Coh. | 105772 | 5.11 | 185217 | 0.09 | 9 | 4.29 |
| CA1 Theta Freq. (Hz) | 308329 | 6.99 | 574674 | 1.13 | 9 | 16.36 |
| CA3 Theta Freq. (Hz) | 296280 | 7.11 | 416176 | 2.68 | 9 | 10.69 |
| DG Theta Freq. (Hz) | 297566 | 11.76 | 432042 | 2.68 | 9 | 6.64 |

F test of the LMM for treatment effects, likelihood ratio test for genotype-treatment interaction effects.

**Table S3.** Differences between genotypes during vehicle treatment. Related to Figures 2–5 and S2–S4.

| Rest Features | PV mean<br>(n = 16) | SST mean<br>(n = 16) | df | F | p |
| --- | --- | --- | --- | --- | --- |
| CA1 MUA during SWRs (Z-score) | 2.64 | 1.44 | 138109 | 2.2 | 0.14 |
| CA3 MUA during SWRs (Z-score) | 1.89 | 0.88 | 138261 | 4.1 | 0.042 |
| DG MUA during SWRs (Z-score) | 1.74 | 0.51 | 135839 | 0.082 | 0.77 |
| SWR Rate (Hz) | 0.45 | 0.44 | 4757 | 1.5 | 0.22 |
| % SWR in Chains | 0.2 | 0.21 | 137661 | 0.26 | 0.61 |
| SWR Length (ms) | 102.99 | 101.34 | 137661 | 0.0073 | 0.93 |
| SWR Frequency (Hz) | 156.24 | 156.15 | 137661 | 0.0004 | 0.98 |
| SW Amplitude (uV) | 375.21 | 384.95 | 471767 | 0.13 | 0.72 |
| SWR CA1 SG Power (Z-score) | 1.31 | 1.25 | 471767 | 0.045 | 0.83 |
| SWR CA1–CA3 Delta Coh. | 0.013 | 0.011 | 1189551 | 0.19 | 0.66 |
| SWR CA1–DG Delta Coh. | 0.011 | 0.0088 | 976070 | 2.4 | 0.12 |
| CA3 SWR Frequency (Hz) | 153.12 | 154.37 | 38901 | 0.69 | 0.41 |
| SWR Size (SD) | 5.93 | 5.93 | 137661 | 0.052 | 0.82 |
| SWR CA3 SG Power (Z-score) | 0.91 | 0.85 | 473237 | 3 | 0.085 |
| SWR DG SG Power (Z-score) | 0.65 | 0.67 | 359153 | 0.29 | 0.59 |
| Run Features | PV mean<br>(n = 8) | SST mean<br>(n = 7) | df | F | p |
| CA1 MUA (Hz) | 87.61 | 136.28 | 155440 | 2.38 | 0.12 |
| CA3 MUA (Hz) | 283.88 | 216.70 | 155440 | 1.79 | 0.18 |
| DG MUA (Hz) | 294.51 | 296.34 | 155400 | 0.00 | 0.99 |
| CA1 SG Power ($\mu V^2$) | 512.08 | 400.01 | 28766 | 0.77 | 0.38 |
| CA1–CA3 SG Coh. | 0.6 | 0.6 | 100543 | $4.20 \times 10^{-5}$ | 0.99 |
| CA1–DG SG Coh. | 0.59 | 0.58 | 114500 | 0.04 | 0.84 |
| CA1 SG Freq. (Hz) | 40.24 | 41.2 | 1066217 | 4.6 | 0.032 |
| CA1 Theta–SG Comod. | 0.0051 | 0.0049 | 28766 | 0.0015 | 0.97 |
| CA1 FG Power ($\mu V^2$) | 79.36 | 63.72 | 28766 | 0.34 | 0.56 |
| CA1 SG/FG ratio | 6.35 | 5.71 | 28766 | 1 | 0.32 |
| CA1–CA3 FG Coh. | 0.54 | 0.56 | 110360 | 0.81 | 0.37 |
| CA1–DG FG Coh. | 0.52 | 0.53 | 95023 | 0.31 | 0.58 |
| CA1 FG Freq (Hz) | 79.7 | 78.93 | 2095627 | 1.3 | 0.25 |
| CA1 Theta–FG Comod. | 0.0079 | 0.0074 | 28766 | 0.036 | 0.85 |
| CA3 SG Power ($\mu V^2$) | 115.94 | 134.21 | 31822 | 1.2 | 0.28 |
| DG SG Power ($\mu V^2$) | 313.19 | 303.08 | 31169 | 0.11 | 0.74 |
| **CA3 SG Freq. (Hz)** | **40.54** | **41.9** | **1089935** | **12** | **0.00042** |
| **DG SG Freq. (Hz)** | **41.06** | **42.4** | **1241681** | **13** | **0.00025** |
| CA3 Theta–SG Comod. | 0.0049 | 0.0053 | 31822 | 2.2 | 0.14 |
| DG Theta–SG Comod. | 0.0047 | 0.0053 | 31169 | 4.5 | 0.034 |
| CA3 FG Power ($\mu V^2$) | 18.3 | 27.1 | 31822 | 5.6 | 0.018 |
| DG FG Power ($\mu V^2$) | 64.54 | 73.9 | 31169 | 2.3 | 0.13 |
| **CA3 SG/FG ratio** | **6.2** | **3.99** | **31822** | **11** | **0.00078** |
| **DG SG/FG ratio** | **5.16** | **3.4** | **31169** | **11** | **0.00071** |
| CA3 FG Freq. (Hz) | 81.99 | 80.18 | 2155990 | 5.2 | 0.022 |
| DG FG Freq. (Hz) | 81.75 | 80.42 | 2381526 | 6.7 | 0.0095 |
| CA3 Theta–FG Comod. | 0.007 | 0.0085 | 31822 | 4 | 0.046 |
| DG Theta–FG Comod. | 0.0086 | 0.009 | 31169 | 0.94 | 0.33 |
| CA1 Theta Power ($\mu V^2$) | 811.3 | 613.97 | 28766 | 0.21 | 0.64 |
| CA3 Theta Power ($\mu V^2$) | 878.78 | 1090.14 | 31822 | 2.3 | 0.13 |
| DG Theta Power ($\mu V^2$) | 2036.8 | 1927.36 | 31169 | 0.35 | 0.56 |
| CA1–CA3 Theta Coh. | 0.81 | 0.8 | 95023 | 0.046 | 0.83 |
| CA1–DG Theta Coh. | 0.79 | 0.77 | 110360 | 0.39 | 0.53 |
| CA1 Theta Freq. (Hz) | 9.18 | 9.09 | 312211 | 0.34 | 0.56 |
| CA3 Theta Freq. (Hz) | 9.15 | 9.03 | 246172 | 0.38 | 0.54 |
| DG Theta Freq. (Hz) | 9.24 | 9.21 | 270460 | 0.056 | 0.81 |

F test of the LMM for genotype effects. Significant comparisons are bolded.

**Table S4.** Statistical details for Figures S5 and S6.

| Rest Features | PV-Cre/ SST-Cre (n = 13) |  | PV-Cre vs PV-Cre/ SST-Cre |  | SST-Cre vs PV-Cre/ SST-Cre |  | Empty Vector (n = 8) |  |
| --- | --- | --- | --- | --- | --- | --- | --- | --- |
| | df | F | df | $\chi^2$ | df | $\chi^2$ | df | F |
| MUA during SWRs (Z-score) | 76435 | 7.16 | 9 | -68724 | 9 | -54461 | 21480 | 0.02 |
| MUA during SWRs (Z-score) | 76435 | 7.78 | 9 | -49460 | 9 | -19733 | 21480 | 0.19 |
| MUA during SWRs (Z-score) | 76435 | 24.13 | 9 | -120845 | 9 | -6232 | 21480 | 1.42 |
| SWR Rate (Hz) | 2709 | 12.79 | 9 | 4.51 | 9 | 19.54 | 892 | 0.11 |
| % SWR in Chains | 75927 | 18.82 | 9 | 1.45 | 9 | 5.19 | 21310 | 0.08 |
| SWR Length (ms) | 75927 | 0.48 | 9 | 5.36 | 9 | 15.06 | 21310 | 0.17 |
| SWR Frequency (Hz) | 75927 | 21.2 | 9 | 12.28 | 9 | 26.56 | 21310 | 0.25 |
| SW Amplitude (uV) | 255972 | 29.64 | 9 | 16.03 | 9 | 29.41 | 66351 | 0.52 |
| CA1-sr SG Power (Z-score) | 255972 | 46.62 | 9 | -26019 | 9 | 28.18 | 66351 | 2.33 |
| SWR Delta SG Coherence | 750476 | 0.99 | 9 | -40005 | 9 | -35500 | 168587 | 0.02 |
| SWR Delta SG Coherence | 524754 | 1.44 | 9 | -97130 | 9 | -63661 | 142513 | 2.35 |
| CA3 SWR Frequency (Hz) | 21295 | 16.73 | 9 | 4.19 | 9 | 8.92 | 8004 | 0.4 |
| SWR Size (SD) | 75927 | 20.15 | 9 | 7.7 | 9 | 18.37 | 21310 | 0.49 |
| SG Power (Z-score) | 303136 | 23.76 | 9 | -15146 | 9 | -26292 | 84625 | 3.48 |
| SG Power (Z-score) | 209470 | 14.65 | 9 | -5617 | 9 | -291 | 68253 | 1.5 |
| Run Features | PV-Cre/ SST-Cre (n = 7) |  | PV-Cre vs PV-Cre/ SST-Cre |  | SST-Cre vs PV-Cre/ SST-Cre |  | Empty Vector (n = 8) |  |
| | df | F | df | $\chi^2$ | df | $\chi^2$ | df | F |
| CA1 MUA (Hz) | 140065 | 4.54 | 9 | 2.69 | 9 | 13.32 | 120735 | 1.44 |
| CA3 MUA (Hz) | 140065 | 37.10 | 9 | 8.28 | 9 | 4.24 | 120735 | 0.85 |
| DG MUA (Hz) | 140065 | 36.95 | 9 | 9.11 | 9 | 18.41 | 120735 | 0.27 |
| CA1 SG Power ($\mu V^2$) | 50966 | 11.62 | 9 | 5.56 | 9 | 6.07 | 45384 | 3.36 |
| CA1–CA3 SG Coh. | 204656 | 11.84 | 9 | 7.35 | 9 | 8.89 | 178143 | 2.26 |
| CA1–DG SG Coh. | 170411 | 10.86 | 9 | 3.28 | 9 | 11.26 | 167832 | 1.06 |
| CA1 SG Freq. (Hz) | 1817692 | 15.1 | 9 | 3.34 | 9 | 17.93 | 1649935 | 1.14 |
| CA1 Theta–SG Comod. | 50966 | 24.58 | 9 | 6.73 | 9 | 3.76 | 45384 | 0.02 |
| CA1 FG Power ($\mu V^2$) | 50966 | 23.04 | 9 | 16.9 | 9 | 1.37 | 49047 | 0.13 |
| CA1 SG/FG ratio | 50966 | 21.37 | 9 | 8.33 | 9 | 28.28 | 49047 | 0.01 |
| CA1–CA3 FG Coh. | 170411 | 0.17 | 9 | 7.95 | 9 | 14.52 | 173082 | 2.64 |
| CA1–DG FG Coh. | 204656 | 0.17 | 9 | 13.66 | 9 | 12.98 | 185143 | 0.05 |
| CA1 FG Freq (Hz) | 3744547 | 14.73 | 9 | 8.25 | 9 | 23.3 | 3333362 | 0.61 |
| CA1 Theta–FG Comod. | 50966 | 13.94 | 9 | 7.99 | 9 | 32.4 | 45384 | 0.01 |
| CA3 SG Power ($\mu V^2$) | 58724 | 0.04 | 9 | 8.69 | 9 | 8.4 | 59762 | 2.92 |
| DG SG Power ($\mu V^2$) | 48576 | 5.71 | 9 | 9.99 | 9 | 6.48 | 51661 | 0.24 |
| CA3 SG Freq. (Hz) | 2137020 | 8.89 | 9 | 6.36 | 9 | 21.1 | 2220964 | 1 |
| DG SG Freq. (Hz) | 1790505 | 8.08 | 9 | 5.33 | 9 | 29.44 | 1961970 | 1.2 |
| CA3 Theta–SG Comod. | 58724 | 12.75 | 9 | 34.51 | 9 | 23.71 | 59762 | 0.23 |
| DG Theta–SG Comod. | 48576 | 42.17 | 9 | 32.26 | 9 | 10.98 | 51661 | 0 |
| CA3 FG Power ($\mu V^2$) | 58724 | 22.57 | 9 | 17.82 | 9 | 31.67 | 59762 | 0.86 |
| DG FG Power ($\mu V^2$) | 48576 | 41.75 | 9 | 19.95 | 9 | 32.09 | 51661 | 0.21 |
| CA3 SG/FG ratio | 58724 | 0.74 | 9 | 8.14 | 9 | 19.44 | 59762 | 1.42 |
| DG SG/FG ratio | 48576 | 1.11 | 9 | 6.75 | 9 | 21.1 | 51661 | 2.11 |
| CA3 FG Freq. (Hz) | 4425768 | 10.28 | 9 | 9.55 | 9 | 14.96 | 4511384 | 0.43 |
| DG FG Freq. (Hz) | 3665938 | 15.43 | 9 | 16.3 | 9 | 22 | 3911662 | 0.11 |
| CA3 Theta–FG Comod. | 58724 | 9.18 | 9 | 6.54 | 9 | 5.48 | 59762 | 1.48 |
| DG Theta–FG Comod. | 48576 | 15.43 | 9 | 5.86 | 9 | 3.03 | 51661 | 0.88 |
| CA1 Theta Power ($\mu V^2$) | 50966 | 30.84 | 9 | 6.49 | 9 | 8.21 | 49047 | 2.7 |
| CA3 Theta Power ($\mu V^2$) | 58724 | 0.86 | 9 | 24.22 | 9 | 20.06 | 59762 | 0.18 |
| DG Theta Power ($\mu V^2$) | 48576 | 6.03 | 9 | 20.96 | 9 | 18.68 | 51661 | 0.26 |
| CA1–CA3 Theta Coh. | 204656 | 7.44 | 9 | 6.77 | 9 | 1.07 | 185143 | 0.84 |
| CA1–DG Theta Coh. | 170411 | 0.19 | 9 | 4.17 | 9 | 3.98 | 173082 | 0.29 |
| CA1 Theta Freq. (Hz) | 529462 | 32.2 | 9 | 16.59 | 9 | 7.85 | 473424 | 0.17 |
| CA3 Theta Freq. (Hz) | 508905 | 29.47 | 9 | 13.03 | 9 | 4.72 | 513283 | 0.11 |
| DG Theta Freq. (Hz) | 421267 | 46.45 | 9 | 6.52 | 9 | 7.92 | 451191 | 0.06 |

F test of the LMM for treatment effects, likelihood ratio test for genotype-treatment interaction effects.

**Table S5.**
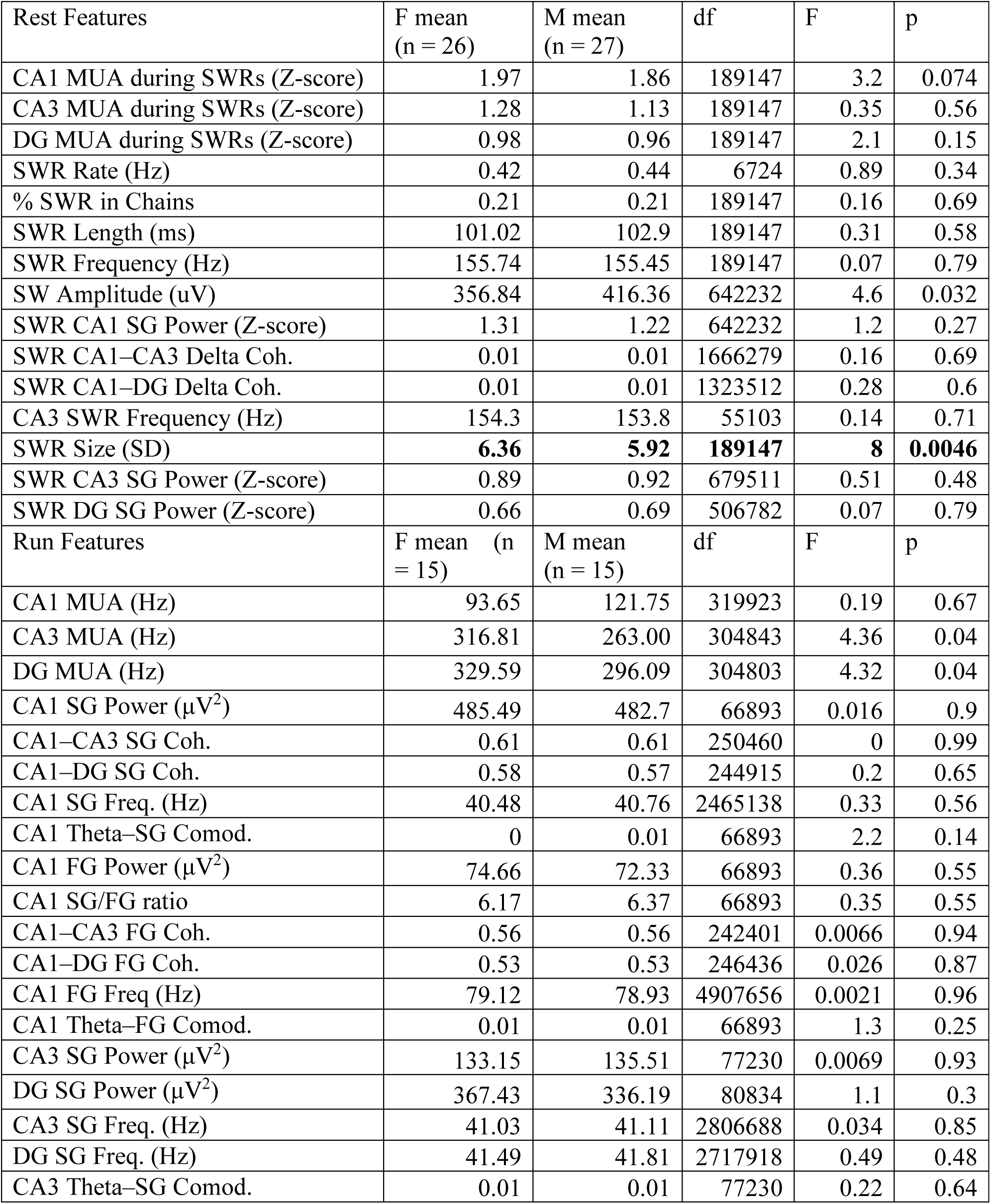
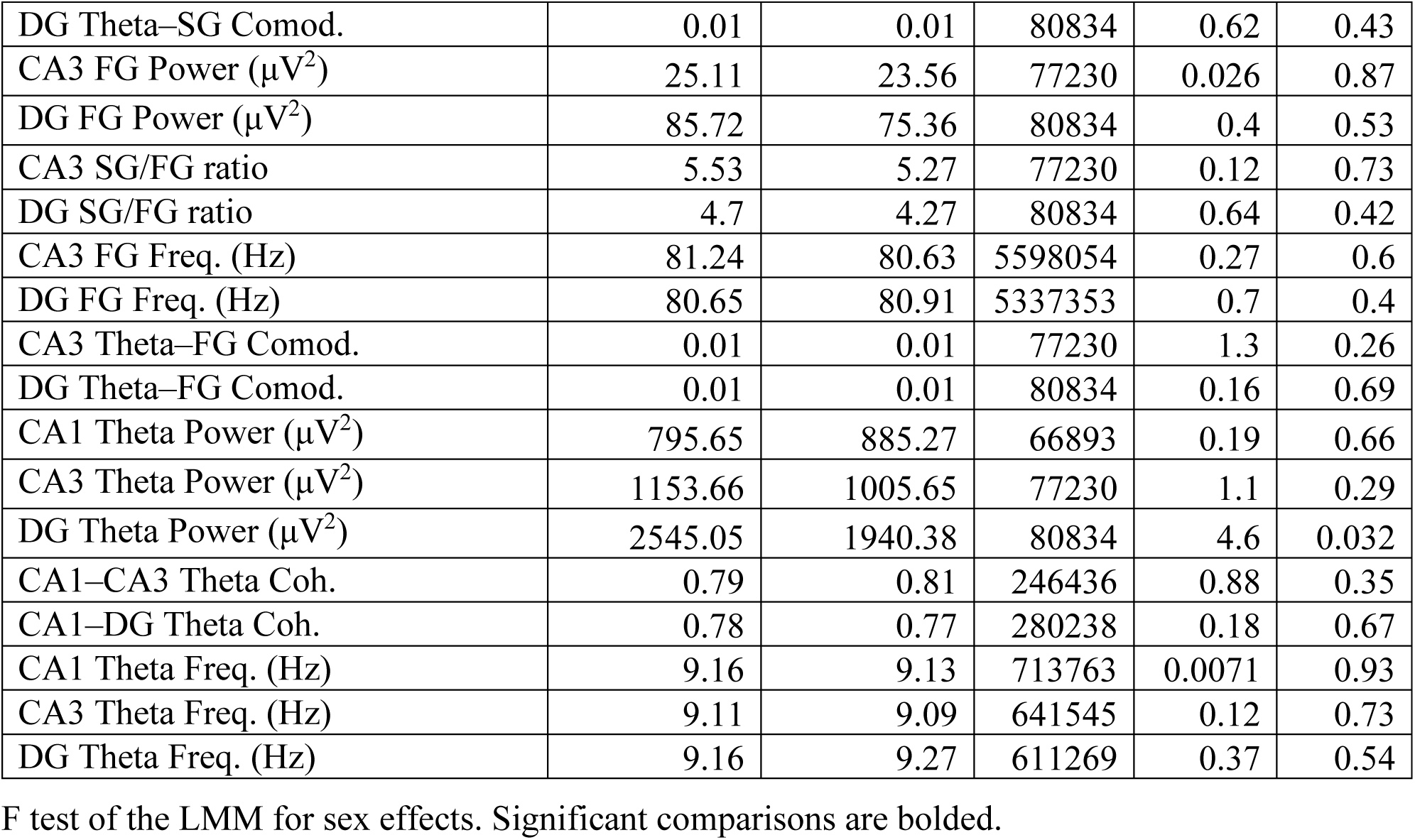
Sex differences across all mice during vehicle treatment. Related to Figures 2–6 and S2–S6.

| Rest Features | F mean<br>(n = 26) | M mean<br>(n = 27) | df | F | p |
| --- | --- | --- | --- | --- | --- |
| CA1 MUA during SWRs (Z-score) | 1.97 | 1.86 | 189147 | 3.2 | 0.074 |
| CA3 MUA during SWRs (Z-score) | 1.28 | 1.13 | 189147 | 0.35 | 0.56 |
| DG MUA during SWRs (Z-score) | 0.98 | 0.96 | 189147 | 2.1 | 0.15 |
| SWR Rate (Hz) | 0.42 | 0.44 | 6724 | 0.89 | 0.34 |
| % SWR in Chains | 0.21 | 0.21 | 189147 | 0.16 | 0.69 |
| SWR Length (ms) | 101.02 | 102.9 | 189147 | 0.31 | 0.58 |
| SWR Frequency (Hz) | 155.74 | 155.45 | 189147 | 0.07 | 0.79 |
| SW Amplitude (uV) | 356.84 | 416.36 | 642232 | 4.6 | 0.032 |
| SWR CA1 SG Power (Z-score) | 1.31 | 1.22 | 642232 | 1.2 | 0.27 |
| SWR CA1–CA3 Delta Coh. | 0.01 | 0.01 | 1666279 | 0.16 | 0.69 |
| SWR CA1–DG Delta Coh. | 0.01 | 0.01 | 1323512 | 0.28 | 0.6 |
| CA3 SWR Frequency (Hz) | 154.3 | 153.8 | 55103 | 0.14 | 0.71 |
| SWR Size (SD) | **6.36** | **5.92** | **189147** | **8** | **0.0046** |
| SWR CA3 SG Power (Z-score) | 0.89 | 0.92 | 679511 | 0.51 | 0.48 |
| SWR DG SG Power (Z-score) | 0.66 | 0.69 | 506782 | 0.07 | 0.79 |
| Run Features | F mean (n = 15) | M mean (n = 15) | df | F | p |
| CA1 MUA (Hz) | 93.65 | 121.75 | 319923 | 0.19 | 0.67 |
| CA3 MUA (Hz) | 316.81 | 263.00 | 304843 | 4.36 | 0.04 |
| DG MUA (Hz) | 329.59 | 296.09 | 304803 | 4.32 | 0.04 |
| CA1 SG Power ($\mu V^2$) | 485.49 | 482.7 | 66893 | 0.016 | 0.9 |
| CA1–CA3 SG Coh. | 0.61 | 0.61 | 250460 | 0 | 0.99 |
| CA1–DG SG Coh. | 0.58 | 0.57 | 244915 | 0.2 | 0.65 |
| CA1 SG Freq. (Hz) | 40.48 | 40.76 | 2465138 | 0.33 | 0.56 |
| CA1 Theta–SG Comod. | 0 | 0.01 | 66893 | 2.2 | 0.14 |
| CA1 FG Power ($\mu V^2$) | 74.66 | 72.33 | 66893 | 0.36 | 0.55 |
| CA1 SG/FG ratio | 6.17 | 6.37 | 66893 | 0.35 | 0.55 |
| CA1–CA3 FG Coh. | 0.56 | 0.56 | 242401 | 0.0066 | 0.94 |
| CA1–DG FG Coh. | 0.53 | 0.53 | 246436 | 0.026 | 0.87 |
| CA1 FG Freq (Hz) | 79.12 | 78.93 | 4907656 | 0.0021 | 0.96 |
| CA1 Theta–FG Comod. | 0.01 | 0.01 | 66893 | 1.3 | 0.25 |
| CA3 SG Power ($\mu V^2$) | 133.15 | 135.51 | 77230 | 0.0069 | 0.93 |
| DG SG Power ($\mu V^2$) | 367.43 | 336.19 | 80834 | 1.1 | 0.3 |
| CA3 SG Freq. (Hz) | 41.03 | 41.11 | 2806688 | 0.034 | 0.85 |
| DG SG Freq. (Hz) | 41.49 | 41.81 | 2717918 | 0.49 | 0.48 |
| CA3 Theta–SG Comod. | 0.01 | 0.01 | 77230 | 0.22 | 0.64 |
| DG Theta–SG Comod. | 0.01 | 0.01 | 80834 | 0.62 | 0.43 |
| CA3 FG Power ($\mu V^2$) | 25.11 | 23.56 | 77230 | 0.026 | 0.87 |
| DG FG Power ($\mu V^2$) | 85.72 | 75.36 | 80834 | 0.4 | 0.53 |
| CA3 SG/FG ratio | 5.53 | 5.27 | 77230 | 0.12 | 0.73 |
| DG SG/FG ratio | 4.7 | 4.27 | 80834 | 0.64 | 0.42 |
| CA3 FG Freq. (Hz) | 81.24 | 80.63 | 5598054 | 0.27 | 0.6 |
| DG FG Freq. (Hz) | 80.65 | 80.91 | 5337353 | 0.7 | 0.4 |
| CA3 Theta–FG Comod. | 0.01 | 0.01 | 77230 | 1.3 | 0.26 |
| DG Theta–FG Comod. | 0.01 | 0.01 | 80834 | 0.16 | 0.69 |
| CA1 Theta Power ($\mu V^2$) | 795.65 | 885.27 | 66893 | 0.19 | 0.66 |
| CA3 Theta Power ($\mu V^2$) | 1153.66 | 1005.65 | 77230 | 1.1 | 0.29 |
| DG Theta Power ($\mu V^2$) | 2545.05 | 1940.38 | 80834 | 4.6 | 0.032 |
| CA1–CA3 Theta Coh. | 0.79 | 0.81 | 246436 | 0.88 | 0.35 |
| CA1–DG Theta Coh. | 0.78 | 0.77 | 280238 | 0.18 | 0.67 |
| CA1 Theta Freq. (Hz) | 9.16 | 9.13 | 713763 | 0.0071 | 0.93 |
| CA3 Theta Freq. (Hz) | 9.11 | 9.09 | 641545 | 0.12 | 0.73 |
| DG Theta Freq. (Hz) | 9.16 | 9.27 | 611269 | 0.37 | 0.54 |

F test of the LMM for sex effects. Significant comparisons are bolded.

**Table S6.** Effects of PV^+^ and/or SST^+^ interneuron suppression stratified by sex. Related to Figures 2–6 and S2–S6.

| PV-Cre: Rest Features | F $\beta$ (n = 9) | M $\beta$ (n = 7) | df | $\chi^2$ | p |
| --- | --- | --- | --- | --- | --- |
| CA1 MUA during SWRs (Z-score) | 0.16 | 0.18 | 9 | -8406 | 1 |
| CA3 MUA during SWRs (Z-score) | 0.14 | 0.27 | 9 | -46879 | 1 |
| DG MUA during SWRs (Z-score) | 0.11 | 0.11 | 9 | -10142 | 1 |
| SWR Rate (Hz) | 0.058 | 0.073 | 9 | 2.3 | 0.99 |
| % SWR in Chains | -0.00065 | -0.00088 | 9 | 10 | 0.33 |
| SWR Length (ms) | -4 | -1.1 | 9 | 2.4 | 0.98 |
| SWR Frequency (Hz) | 2.3 | 3 | 9 | 1.4 | 1 |
| SW Amplitude (uV) | 37 | 37 | 9 | 2 | 0.99 |
| SWR CA1 SG Power (Z-score) | 0.66 | 0.72 | 9 | -20193 | 1 |
| SWR CA1–CA3 Delta Coh. | 0.013 | 0.0089 | 9 | -47795 | 1 |
| SWR CA1–DG Delta Coh. | 0.015 | 0.0038 | 9 | -29872 | 1 |
| CA3 SWR Frequency (Hz) | -1.1 | -0.67 | 9 | 5.7 | 0.77 |
| SWR Size (SD) | -0.29 | -0.33 | 9 | 3.2 | 0.96 |
| SWR CA3 SG Power (Z-score) | 0.23 | 0.35 | 9 | -21508 | 1 |
| SWR DG SG Power (Z-score) | 0.14 | 0.3 | 9 | -5520 | 1 |
| PV-Cre: Run Features | F $\beta$ (n = 4) | M $\beta$ (n = 4) | df | $\chi^2$ | p |
| CA1 MUA (Hz) | 1.23 | -0.07 | 9 | 12 | 0.19 |
| **CA3 MUA (Hz)** | **367.64** | **3.33** | **9** | **40** | **9.2x10<sup>-6</sup>** |
| **DG MUA (Hz)** | **441.12** | **4.24** | **9** | **53** | **2.6x10<sup>-8</sup>** |
| CA1 SG Power ($\mu V^2$) | 458 | 156 | 9 | 5 | 0.81 |
| CA1–CA3 SG Coh. | 0.076 | 0.015 | 9 | 12 | 0.21 |
| CA1–DG SG Coh. | 0.06 | -0.0086 | 9 | 25 | 0.0032 |
| CA1 SG Freq. (Hz) | -1.4 | -1.4 | 9 | 19 | 0.028 |
| CA1 Theta–SG Comod. | 0.000063 | -0.00059 | 9 | 12 | 0.2 |
| CA1 FG Power ($\mu V^2$) | 24 | 8.9 | 9 | 9 | 0.47 |
| CA1 SG/FG ratio | 4.9 | 3 | 9 | 4 | 0.92 |
| CA1–CA3 FG Coh. | -0.022 | 0.017 | 9 | 20 | 0.017 |
| CA1–DG FG Coh. | -0.044 | 0.011 | 9 | 17 | 0.047 |
| CA1 FG Freq (Hz) | 0.33 | 1.3 | 9 | 5 | 0.82 |
| CA1 Theta–FG Comod. | -0.00031 | -0.0011 | 9 | 7 | 0.61 |
| CA3 SG Power ($\mu V^2$) | 45 | 8.9 | 9 | 6 | 0.76 |
| DG SG Power ($\mu V^2$) | 50 | 2.8 | 9 | 7 | 0.6 |
| CA3 SG Freq. (Hz) | -1.4 | -1.1 | 9 | 8 | 0.52 |
| DG SG Freq. (Hz) | -1.4 | -0.79 | 9 | 8 | 0.57 |
| CA3 Theta–SG Comod. | 0.000041 | 0.0004 | 9 | 11 | 0.3 |
| DG Theta–SG Comod. | 0.00026 | 0.00047 | 9 | 6 | 0.78 |
| CA3 FG Power ($\mu V^2$) | 5 | -0.66 | 9 | 13 | 0.16 |
| DG FG Power ($\mu V^2$) | 1.5 | -5.5 | 9 | 5 | 0.79 |
| CA3 SG/FG ratio | 1.8 | 0.74 | 9 | 11 | 0.3 |
| DG SG/FG ratio | 1.4 | 0.52 | 9 | 9 | 0.46 |
| CA3 FG Freq. (Hz) | 0.19 | 1.9 | 9 | 17 | 0.053 |
| DG FG Freq. (Hz) | 1 | 0.3 | 9 | 10 | 0.35 |
| CA3 Theta-FG Comod. | -0.00016 | -0.00091 | 9 | 9 | 0.4 |
| DG Theta-FG Comod. | -0.0011 | -0.0017 | 9 | 5 | 0.81 |
| CA1 Theta Power ($\mu V^2$) | 1318 | 202 | 9 | 14 | 0.14 |
| CA3 Theta Power ($\mu V^2$) | 143 | -147 | 9 | 5 | 0.79 |
| DG Theta Power ($\mu V^2$) | -104 | -380 | 9 | 9 | 0.48 |
| CA1-CA3 Theta Coh. | 0.033 | -0.068 | 9 | 5 | 0.81 |
| CA1-DG Theta Coh. | -0.019 | -0.12 | 9 | 4 | 0.92 |
| CA1 Theta Freq. (Hz) | 0.047 | 0.43 | 9 | 17 | 0.049 |
| CA3 Theta Freq. (Hz) | 0.075 | 0.5 | 9 | 24 | 0.0036 |
| DG Theta Freq. (Hz) | 0.15 | 0.53 | 9 | 25 | 0.0034 |
| SST-Cre: Rest Features | F $\beta$ (n = 5) | M $\beta$ (n = 11) | df | $\chi^2$ | p |
| CA1 MUA during SWRs (Z-score) | -0.14 | 0.042 | 9 | -57534 | 1 |
| CA3 MUA during SWRs (Z-score) | 0.16 | 0.2 | 9 | -3606 | 1 |
| DG MUA during SWRs (Z-score) | 0.12 | 0.17 | 9 | -81019 | 1 |
| SWR Rate (Hz) | -0.049 | -0.036 | 9 | 2 | 0.99 |
| % SWR in Chains | -0.0015 | -0.00046 | 9 | 10 | 0.32 |
| SWR Length (ms) | 11 | 4.8 | 9 | 12 | 0.24 |
| SWR Frequency (Hz) | -4.3 | -0.78 | 9 | 22 | 0.0093 |
| **SW Amplitude ($\mu V$)** | **-5.8** | **14** | **9** | **27** | **0.0016** |
| SWR CA1 SG Power (Z-score) | -0.092 | 0.19 | 9 | -32461 | 1 |
| SWR CA1-CA3 Delta Coh. | -0.012 | -0.00035 | 9 | -20181 | 1 |
| SWR CA1-DG Delta Coh. | -0.0042 | -0.00094 | 9 | -83276 | 1 |
| CA3 SWR Frequency (Hz) | -1.3 | 0.05 | 9 | 4.1 | 0.9 |
| SWR Size (SD) | 0.091 | 0.24 | 9 | 6.6 | 0.68 |
| SWR CA3 SG Power (Z-score) | -0.027 | 0.19 | 9 | -15984 | 1 |
| SWR DG SG Power (Z-score) | 0.051 | 0.19 | 9 | -28517 | 1 |
| SST-Cre: Run Features | F $\beta$ (n = 3) | M $\beta$ (n = 4) | df | $\chi^2$ | p |
| CA1 MUA (Hz) | -18.59 | 1.42 | 9 | 10 | 0.36 |
| **CA3 MUA (Hz)** | **308.17** | **2.33** | **9** | **32** | **0.00021** |
| **DG MUA (Hz)** | **437.16** | **2.59** | **9** | **39** | **1.3x10<sup>-5</sup>** |
| CA1 SG Power ($\mu V^2$) | 53 | 51 | 9 | 11 | 0.27 |
| CA1-CA3 SG Coh. | -0.034 | 0.015 | 9 | 8 | 0.57 |
| CA1-DG SG Coh. | -0.027 | 0.00079 | 9 | 13 | 0.14 |
| CA1 SG Freq. (Hz) | 0.34 | -0.41 | 9 | 17 | 0.052 |
| CA1 Theta-SG Comod. | -1.30x10<sup>-4</sup> | -0.00037 | 9 | 21 | 0.012 |
| **CA1 FG Power ($\mu V^2$)** | **36** | **8.4** | **9** | **34** | **0.000086** |
| CA1 SG/FG ratio | -0.8 | -1.2 | 9 | 13 | 0.18 |
| CA1–CA3 FG Coh. | 0.003 | 0.035 | 9 | 15 | 0.086 |
| CA1–DG FG Coh. | 0.013 | 0.026 | 9 | 20 | 0.016 |
| CA1 FG Freq. (Hz) | 1.6 | 2.5 | 9 | 14 | 0.13 |
| CA1 Theta–FG Comod. | 0.00076 | 0.00068 | 9 | 11 | 0.25 |
| CA3 SG Power ($\mu V^2$) | -12 | -36 | 9 | 18 | 0.03 |
| DG SG Power ($\mu V^2$) | -40 | -73 | 9 | 11 | 0.28 |
| CA3 SG Freq. (Hz) | 0.17 | -0.2 | 9 | 23 | 0.0052 |
| DG SG Freq. (Hz) | 0.0015 | -0.15 | 9 | 17 | 0.055 |
| CA3 Theta–SG Comod. | $-1.30 \times 10^{-4}$ | -0.00016 | 9 | 17 | 0.046 |
| DG Theta–SG Comod. | $-7.10 \times 10^{-5}$ | -0.00021 | 9 | 5 | 0.87 |
| CA3 FG Power ($\mu V^2$) | 2.5 | 5.6 | 9 | 15 | 0.1 |
| DG FG Power ($\mu V^2$) | 17 | 16 | 9 | 9 | 0.45 |
| CA3 SG/FG ratio | -0.062 | -0.52 | 9 | 23 | 0.0053 |
| DG SG/FG ratio | -0.1 | -0.32 | 9 | 13 | 0.17 |
| CA3 FG Freq. (Hz) | 1.5 | 2.8 | 9 | 18 | 0.033 |
| DG FG Freq. (Hz) | 1.6 | 2.9 | 9 | 25 | 0.003 |
| CA3 Theta–FG Comod. | -0.00036 | -0.00022 | 9 | 12 | 0.22 |
| DG Theta–FG Comod. | -0.00083 | -0.00049 | 9 | 7 | 0.67 |
| CA1 Theta Power ($\mu V^2$) | 712 | 204 | 9 | 16 | 0.061 |
| CA3 Theta Power ($\mu V^2$) | 65 | -372 | 9 | 6 | 0.73 |
| DG Theta Power ($\mu V^2$) | -114 | -633 | 9 | 5 | 0.83 |
| CA1–CA3 Theta Coh. | 0.043 | 0.028 | 9 | 5 | 0.84 |
| CA1–DG Theta Coh. | -0.013 | 0.028 | 9 | 11 | 0.25 |
| CA1 Theta Freq. (Hz) | -0.1 | 0.25 | 9 | 22 | 0.0086 |
| CA3 Theta Freq. (Hz) | -0.027 | 0.31 | 9 | 20 | 0.017 |
| DG Theta Freq. (Hz) | 0.034 | 0.28 | 9 | 10 | 0.38 |
| PV-Cre/SST-Cre: Rest Features | F $\beta$ (n = 8) | M $\beta$ (n = 5) | df | $\chi^2$ | p |
| CA1 MUA during SWRs (Z-score) | 0.56 | 0.23 | 9 | -10100 | 1 |
| CA3 MUA during SWRs (Z-score) | 0.24 | 0.25 | 9 | -5390 | 1 |
| DG MUA during SWRs (Z-score) | 0.23 | 0.21 | 9 | -33509 | 1 |
| SWR Rate (Hz) | 0.046 | 0.056 | 9 | 2.3 | 0.98 |
| % SWR in Chains | -0.00085 | -0.0013 | 9 | 6.2 | 0.72 |
| SWR Length (ms) | 0.21 | -3.9 | 9 | 4.7 | 0.86 |
| SWR Frequency (Hz) | 3.8 | 1.7 | 9 | 10 | 0.35 |
| SW Amplitude ($\mu V$) | 47 | 37 | 9 | 21 | 0.012 |
| SWR CA1 SG Power (Z-score) | 0.99 | 0.68 | 9 | -2088 | 1 |
| SWR CA1–CA3 Delta Coh. | -0.0017 | -0.0086 | 9 | -43139 | 1 |
| SWR CA1–DG Delta Coh. | 0.0046 | -0.00038 | 9 | -30365 | 1 |
| CA3 SWR Frequency (Hz) | -2 | -1.4 | 9 | 14 | 0.14 |
| SWR Size (SD) | -0.53 | -0.6 | 9 | 11 | 0.24 |
| SWR CA3 SG Power (Z-score) | 0.57 | 0.3 | 9 | -936 | 1 |
| SWR DG SG Power (Z-score) | 0.43 | 0.17 | 9 | -7482 | 1 |
| PV-Cre/SST-Cre: Run Features | F $\beta$ (n = 4) | M $\beta$ (n = 3) | df | $\chi^2$ | p |
| CA1 MUA (Hz) | 11.02 | 0.13 | 9 | 16.26 | 0.06 |
| **CA3 MUA (Hz)** | **335.91** | **2.59** | **9** | **23.31** | **0.0055** |
| **DG MUA (Hz)** | **336.24** | **2.71** | **9** | **23.12** | **0.0059** |
| CA1 SG Power ($\mu V^2$) | 180 | 24 | 9 | 15 | 0.079 |
| CA1–CA3 SG Coh. | 0.032 | 0.0097 | 9 | 17 | 0.056 |
| CA1–DG SG Coh. | 0.031 | 0.11 | 9 | 22 | 0.011 |
| CA1 SG Freq. (Hz) | -2 | -1.4 | 9 | 17 | 0.051 |
| CA1 Theta–SG Comod. | -0.00037 | -0.00025 | 9 | 9.7 | 0.38 |
| CA1 FG Power ($\mu V^2$) | 28 | 30 | 9 | 18 | 0.033 |
| CA1 SG/FG ratio | 2.6 | 1.4 | 9 | 14 | 0.11 |
| CA1–CA3 FG Coh. | 0.0051 | 0.0009 | 9 | 13 | 0.18 |
| CA1–DG FG Coh. | 0.0012 | 0.0085 | 9 | 14 | 0.11 |
| CA1 FG Freq. (Hz) | 3.1 | 1.3 | 9 | 17 | 0.046 |
| CA1 Theta–FG Comod. | -0.001 | -0.00061 | 9 | 9 | 0.44 |
| CA3 SG Power ($\mu V^2$) | -5.8 | 12 | 9 | 17 | 0.043 |
| DG SG Power ($\mu V^2$) | -66 | 1.3 | 9 | 15 | 0.088 |
| CA3 SG Freq. (Hz) | -2.1 | -1.3 | 9 | 17 | 0.043 |
| DG SG Freq. (Hz) | -1.7 | -1.1 | 9 | 17 | 0.052 |
| CA3 Theta–SG Comod. | -0.00042 | -0.00019 | 9 | 6 | 0.74 |
| DG Theta–SG Comod. | -0.00054 | -0.00031 | 9 | 14 | 0.11 |
| CA3 FG Power ($\mu V^2$) | -2.5 | -1.9 | 9 | 13 | 0.15 |
| DG FG Power ($\mu V^2$) | -20 | -12 | 9 | 14 | 0.11 |
| CA3 SG/FG ratio | 0.33 | 0.88 | 9 | 17 | 0.053 |
| DG SG/FG ratio | 0.28 | 0.45 | 9 | 16 | 0.07 |
| CA3 FG Freq. (Hz) | 3 | 0.93 | 9 | 17 | 0.049 |
| DG FG Freq. (Hz) | 3.3 | 1.4 | 9 | 15 | 0.089 |
| CA3 Theta–FG Comod. | -0.00054 | -0.00096 | 9 | 11 | 0.25 |
| DG Theta–FG Comod. | -0.0011 | -0.0023 | 9 | 13 | 0.17 |
| CA1 Theta Power ($\mu V^2$) | 1021 | 191 | 9 | 24 | 0.0051 |
| CA3 Theta Power ($\mu V^2$) | -20 | -332 | 9 | 17 | 0.052 |
| DG Theta Power ($\mu V^2$) | -332 | -951 | 9 | 21 | 0.012 |
| CA1–CA3 Theta Coh. | 0.06 | -0.011 | 9 | 17 | 0.041 |
| CA1–DG Theta Coh. | 0.0093 | -0.13 | 9 | 19 | 0.026 |
| CA1 Theta Freq. (Hz) | 0.36 | 0.23 | 9 | 13 | 0.17 |
| CA3 Theta Freq. (Hz) | 0.38 | 0.23 | 9 | 13 | 0.15 |
| DG Theta Freq. (Hz) | 0.39 | 0.33 | 9 | 13 | 0.14 |

835 Likelihood ratio test for sex-treatment interaction effects. Significant comparisons are bolded.

## KEY RESOURCES TABLE

| REAGENT or RESOURCE | SOURCE | IDENTIFIER |
| --- | --- | --- |
| **Antibodies** |  |  |
| Rat anti-SST | Millipore | Cat#MAB354;<br>RRID:AB_2255365 |
| Mouse anti-PV | Millipore | Cat#MAB1572;<br>RRID:AB_2174013 |
| Biotinylated rabbit anti-RFP | Abcam | Cat#ab34771; RRID:AB_777699 |
| Donkey anti-rat AlexaFluor488 | Thermo Fisher Scientific | Cat#A-21208;<br>RRID:AB_2535794 |
| Goat anti-rat AlexaFluor647 | Thermo Fisher Scientific | Cat#A-21247;<br>RRID:AB_141778 |
| Donkey anti-mouse<br>AlexaFluor647 | Thermo Fisher Scientific | Cat#A-31571;<br>RRID:AB_162542 |
| Streptavidin AlexaFluor594 | Sigma Aldrich | Cat#S6402 |
| **Bacterial and Virus Strains** |  |  |
| AAV5-hSyn-DIO-hM4D(Gi)-<br>mCherry | UNC Viral Vector Core<br>(Krashes et al., 2011) | N/A |
| AAV5-hSyn-DIO-mCherry | Addgene | RRID:Addgene_50459 |
| **Chemicals, Peptides, and Recombinant Proteins** |  |  |
| Ketamine | Henry Schein | Cat#1049007 |
| Xylazine (Anased) | Henry Schein | Cat#1311139 |
| Isoflurane | Henry Schein | Cat#029405 |
| Buprenorphine | Henry Schein | Cat#055175 |
| Ketofen | Henry Schein | Cat#005487 |
| Avertin (2,2,2-Tribromoethanol) | Millipore Sigma | Cat# T48402 |
| Dental Adhesive (Metabond) | Parkell | Cat#S396, S398 and S371 |
| Dental Acrylic | Stoelting Co. | Cat#51459 |
| Cresyl Violet Acetate | Sigma Aldrich | Cat#C5042 |
| ProLong Gold with DAPI | Thermo Fisher Scientific | Cat#P36931 |
| **Deposited Data** |  |  |
| 0–300 Hz filtered LFP, MUA<br>spike times, mouse position<br>tracking, and metadata for each<br>electrode site, mouse, and<br>recording session | This paper | Will be uploaded after paper<br>acceptance |
| **Experimental Models: Organisms/Strains** |  |  |
| Mouse: PVCre: *Pvalb<sup>tm1(cre)Arbr</sup>/J* | Jackson Laboratory | MGI:3590684;<br>RRID:IMSR_JAX:013044 |
| Mouse: SSTCre: *Sst<sup>tm2.1(cre)Zjh</sup>/J* | Jackson Laboratory | MGI:4838416;<br>RRID:IMSR_JAX:008069 |
| **Software and Algorithms** |  |  |
| Custom scripts for this paper | This paper | Will be uploaded after paper<br>acceptance |
| Trodes | SpikeGadgets | https://bitbucket.org/mkarlssso/trodes/downloads |
| NeuroQuery | Bitbucket | https://bitbucket.org/mkarlssso/neuroquery/src/master |
| MATLAB | Mathworks | N/A |
| Chronux | Cold Spring Harbor Laboratory | http://chronux.org |
| plot spread points | MATLAB Central File Exchange | https://www.mathworks.com/matlabcentral/fileexchange/37105-plot-spread-points-beeswarm-plot |
| Axon pCLAMP | Molecular Devices | https://mdc.custhelp.com/app/answers/detail/a_id/18779/~/axon%E2%84%A2pclamp%E2%84%A2-10-electrophysiology-data-acquisition-%26-analysis-software-download |
| IGOR Pro | WaveMetrics | https://www.wavemetrics.com/ |
| Other |  |  |
| Probe | Neuronexus | A4x8-400-200-704-CM32 |

## STAR METHODS

### Lead Contact and Materials Availability

This study did not generate any unique reagents. Further information and requests for resources and reagents should be directed to and will be fulfilled by the Lead Contacts, Yadong Huang or Loren Frank.

## EXPERIMENTAL MODEL AND SUBJECT DETAILS

C57BL6/J mice with the SST-IRES-Cre allele (*Sst^tm2.1(cre)Zjh^*/J) or the PV-IRES-Cre allele (*Pvalb^tm1(cre)Arbr^*/J) knocked-in (Hippenmeyer et al., 2005; Taniguchi et al., 2011) were originally obtained from Jackson Laboratory. Equal numbers of PV-Cre, SST-Cre and PV-Cre/SST-Cre mice were selected from littermates of a PV-Cre x SST-Cre cross. All animals were bred in-house using trio breeding producing 10 pups per litter on average, which were weaned at 28 days. Equal proportions of males and females aged 3–8 months were selected for each genotype and viral vector. Within each genotype group and sex, mice were randomly assigned to receive either hM4Di-mCherry vector or mCherry empty vector injection. Experimenters were blinded to genotype during surgery and blinded to genotype and viral vector expression during all post-operative behavior, recordings, and histology. Animals were housed in a pathogen-free barrier facility on a 12h light cycle (lights on at 7am and off at 7pm) at 19–23°C and 30–70% humidity. Animals were identified by ear punch under brief isofluorane anesthesia and genotyped by PCR of a tail clipping at both weaning and perfusion. All animals otherwise received no procedures except those reported in this study. Throughout the study, mice were singly housed. All animal experiments were conducted in accordance with the guidelines and regulations of the National Institutes of Health, the University of California, and the Gladstone Institutes under IACUC protocol AN117112.

## METHOD DETAILS

This study consisted of two cohorts. The first cohort (n = 9 PV-Cre, n = 10 SST-Cre, n = 6 PV- Cre/SST-Cre) had home cage recordings (Figure 2) while the second cohort (n = 8 PV-Cre, n = 7 SST-Cre, n = 7 PV-Cre/SST-Cre, and n = 8 empty vector mice) had linear track recordings (Figure 4), then home cage recordings (Figure 2). In the second cohort, 1 PV-Cre and 1 SST-Cre animal died between linear track and home cage recordings and so were not included in the analysis in Figure 2 and Table S1.

### Surgery

Mice were anesthetized by intraperitoneal injection of ketamine (60 mg/kg) and xylazine (30 mg/kg); anesthesia was maintained with 0.6–1.5% isofluorane given through a vaporizer and nose cone. The head was secured with earbars and a tooth bar in a stereotaxic alignment system (Kopf Instruments). Fur was removed from the scalp, which was then sterilized with alternating swabs of chlorhexidine and 70% ethanol. The scalp was opened, sterilized with 3% hydrogen peroxide, and thoroughly cleaned to reduce risk of tissue regrowth. 0.5 mm craniotomies were made at 1.95 mm AP and ± 1.5 mm ML from bregma for viral injection. 1 µL of 4.6 x 10^12^vg/mL AAV5-hSyn-DIO-hM4D(Gi)-mCherry (UNC Viral Vector Core; (Krashes et al., 2011)) or 7 x 10^12^vg/mL AAV5-hSyn-DIO-mCherry (Addgene) was injected at 2.1 mm below the surface of the brain (Andrews-Zwilling et al., 2012; Stefanelli et al., 2016) at an infusion rate of 100nL/min. Skull screws (FST) were inserted into craniotomies over the right frontal cortex and left parietal cortex to anchor and support the implant, and were secured with dental adhesive (C&B Metabond, Parkell). An additional 0.5 mm craniotomy was made over the right cerebellum for insertion of the indifferent ground and reference wires. The craniotomy centered at -1.95 mm AP and 1.5 mm ML from bregma and extended bidirectionally along the ML axis to 2 mm width to receive the recording probe. The probes had four 5 mm shanks spaced 400 µm apart with 8 electrode sites per shank and 200 µm spacing between sites (Neuronexus; configuration A4x8-400-200-704-CM32). The probe was quickly lowered until the tip reached 2.2 mm below the surface of the brain, and the reference and ground wire was inserted into the subdural space above the cerebellum. The probe was cemented in place with dental acrylic and the scalp was closed with nylon sutures. Mice were treated with 0.0375 mg/kg buprenorphine intraperitoneally and 5 mg/kg ketofen subcutaneously 30–45 min after surgery, monitored until ambulatory, then monitored daily for 3 days. A minimum of 3 weeks was allowed for recovery and viral expression before recording.

### Electrophysiology

Each animal was randomly assigned a time during the light cycle, and behavior and recordings were always conducted with that animal at that same time each day ± 1 hour. CNO (NIMH, C-929) in 1% DMSO in 0.9% sterile saline (CNO) or equivalent volume of 1% DMSO in 0.9% sterile saline (vehicle) was administered via intraperitoneal injection at a dose of 2 mg/kg 1 hour prior to data collection. Body weight was measured weekly during treatment and injection volume was adjusted accordingly. Injections were well tolerated and had no adverse effects on health. Data were collected on a Main Control Unit (SpikeGadgets) with simultaneous video tracking at 30 frames/s using Trodes software (SpikeGadgets).

During home cage recordings, data were collected, amplified, multiplexed, processed, and digitized using a 32-channel upright headstage and commutator (SpikeGadgets) at 30 kHz. Data were collected during 60 min home cage sessions for 6 days, alternating vehicle and CNO administration. Home cages were changed to Alpha-dri bedding (Shepherd Specialty Papers) to enable video tracking.

For linear track recordings, animals were screened for sufficient food motivation prior to surgery. Each animal was restricted to 85–90% of its baseline weight then run on a 36 cm linear track for 30 min sessions daily for 10 µL soy milk reward dispensed from custom automatic solenoids controlled by an Environmental Control Unit (SpikeGadgets) with custom scripts. Animals which did not achieve at least 180 pokes over 3 sessions were excluded from surgery. Following surgery, animals ran for 30 min on linear track for 6 days, alternating vehicle and CNO administration. Data were collected, amplified, multiplexed, processed, and digitized using a wireless 32-channel mini logger headstage (SpikeGadgets) at 20 kHz.

### Histology

Mice were deeply anesthetized with avertin, and a 30 µA current was passed through each recording site for 2 s to generate small electrolytic lesions (Ugo Basile). Mice were then perfused with 0.9% NaCl. The brains were removed and stored at 4°C, then fixed in 4% PFA for 2 days, rinsed in PBS for 1 day, and cryoprotected in 30% sucrose for at least 2 days. Brains were cut into 30 µm coronal sections with a microtome (Leica) and stored in cryoprotectant at -20°C. Every third section was stained with cresyl violet, then electrolytic lesion locations were observed under a light microscope (Leica). Every tenth section was used for immunohistochemistry. Sections were blocked and permeabilized in 10% normal donkey serum and 0.5% Triton X for 1 hour at room temperature, and then incubated overnight at 4°C in 1:100 rat anti-SST (Millipore), 1:1000 mouse anti-PV (Millipore), and 1:500 biotinylated rabbit anti-RFP (Abcam). Sections were then incubated for 1 hour at room temperature in 1:1500 donkey anti-rat AlexaFluor488 (Thermo Fisher Scientific) or 1:1500 goat anti-rat AlexaFluor647 (Thermo Fisher Scientific), 1:1500 donkey anti-mouse AlexaFluor647 (Thermo Fisher Scientific), and 1:1000 Streptavidin AlexaFluor594 (Sigma Aldrich) and mounted to slides using DAPI. Images were collected on a fluorescent microscope (Keyence) and counted manually in ImageJ. 1 SST-Cre animal from the first cohort was excluded from all analyses due to low viral expression. Representative images were adjusted for contrast only.

### *Ex vivo* electrophysiology

3–6 month old mice were stereotaxically injected with hM4D as described above, then the scalp was sutured closed. 3 weeks following surgery to allow for viral expression, mice were deeply anesthetized with isofluorane. The brain was rapidly removed and placed in 4°C slicing solution comprised of 110 mM Choline Chloride, 2.5 mM KCl, 1.25 mM NaH_2_PO_4_, 26 mM NaHCO_3_, 2 mM CaCl_2_, 1.3 mM Na Pyruvate, 1 mM L-Ascorbic Acid, and 10 mM dextrose. 300µm sagittal sections were cut using a vibratome (VT 1200s, Leica), transferred to a vapor interface chamber aerated with 95% O_2_ / 5% CO_2_ gas mixture, and allowed to recover at 34°C for one hour prior to recording. Sections were then transferred to a submerged recording chamber at 34°C perfused at 10 mL/min with oxygenated aCSF solution comprised of 124 mM NaCl, 26 mM NaHCO_3_, 10 mM Glucose, 1.25 mM NaH_2_PO_4_, 2.5 mM KCl, 1.25 mM MgCl_2_, and 1.5 mM CaCl_2_. SST^+^ and PV^+^ cells were visually identified in the DG by mCherry expression and morphology using a modified Olympus BXW-51 microscope (Scientifica, Inc). Interneurons were recorded using patch-clamp electrodes filled with an intracellular solution comprised of 125 mM K-gluconate, 10 mM KCl, 10 mM HEPES, 2 mM MgCl_2_, 10 mM EGTA, 4 mM MgATP, 10 mM Na-phosphocreatine, and 3 mM Na_2_GTP. CNO was dissolved to 1μM in aCSF and delivered through the perfusion system. Whole cell recordings were performed using a Multiclamp 700B amplifier (Molecular Devices). The signals were sampled at 10 kHz and digitized using Digidata 1550B with Axon pCLAMP (Molecular Devices). Data was analyzed using custom scripts in IGOR Pro (WaveMetrics).

### Analysis of neural data

Neural data was analyzed with custom software written in MATLAB (Mathworks) with the Chronux toolbox (http://www.chronux.org) and Trodes to MATLAB software (SpikeGadgets). The anatomical location of each electrode site was determined by examining Nissl-stained histological sections, raw LFP traces, the SWR-triggered spectrogram signature, and dentate spikes. Only DG sites with visually confirmed dentate spikes were included in analysis. Data were data were referenced to a corpus callosum electrode, band-pass Butterworth filtered at 0.1–300 Hz, and then downsampled to 1 kHz and analyzed as LFP. All measurements were analyzed per session and electrode site, then averaged across all sessions and all electrode sites within each subregion. Thus, each mouse contributed a single number to all comparisons.

Raw LFP data during rest sessions were band-pass equiripple filtered at 125–200 Hz for SWRs, 0–30 Hz for SWs, and 30–50 Hz for SG. SWRs were detected on the CA1 site closest to the center of the pyramidal layer and defined by the Hilbert envelope of the ripple-filtered trace, smoothed with a 4 ms Gaussian, exceeding 3 SD above baseline for at least 15 ms (Cheng and Frank, 2008). Analysis was restricted to periods of extended immobility, after the mouse Gaussian smoothed velocity had been < 1 cm/s for 30 seconds or more. SWRs were considered part of chains if a second SWR event occurred within 200 ms of the end of an event. Instantaneous frequency was defined by interpeak times during SWRs. SW amplitude was defined as the maximum absolute value of the Hilbert envelope of the SW-filtered trace during SWRs. SWRs in CA3 were detected on the site with the highest MUA and examined only when they coincided with an SWR detected in CA1.

SWR-triggered spectrograms for each electrode site and SWR-triggered coherence between regions were calculated with the multitaper method, as previously described (Carr et al., 2012), with a 100 ms sliding window. Delta coherence was calculated as the difference between the 100 ms window starting 400 ms before SWR onset and the 100 ms window after SWR onset; SWRs that were preceded by SWRs within this window were excluded. For illustration in figures, a 10 ms sliding window was used. SWR-associated SG power was calculated as the averaged z-scored power over the 30–50 Hz frequency band 0–100 ms after ripple detection. SG power was analyzed for three regions: CA1-sr, CA3 including pyr and sr, and DG including hilus and granule cell layers. For CNO epochs, the mean and SD of the SG filtered signal from the vehicle epoch recorded the day before were used for z-scoring.

For MUA analysis, data were referenced to a corpus callosum electrode, band-pass filtered at 600–6000 Hz (Butterworth), and then events greater than 75 μV were treated as spikes. Sites used for SWR detection were further verified to be in the CA1 pyramidal layer as they showed large increases in MUA during SWRs. The site closest to the center of the cell layer, as determined by highest MUA, was used for MUA analysis. For fast ripple analysis, data were downsampled to 5 kHz and band-pass equiripple filtered at 125–600 Hz, then events were detected in CA1 when the Hilbert envelope of the ripple-filtered trace, smoothed with a 4 ms Gaussian, exceeded 3 SD above baseline for at least 3 oscillations of the filtered trace. Events were classified as fast ripples if the mean frequency was above 250 Hz.

Slow and fast gamma bands were defined by the frequencies with highest cross-frequency coupling, as previously described (Colgin et al., 2009; Kemere et al., 2013). Raw LFP during linear track sessions were then band-pass least squares FIR filtered at 5–11 Hz for theta, 20–50 Hz for SG, and 50–110 Hz for FG; these definitions matched those previously defined in mice using time-frequency methods (Cabral et al., 2014; Chen et al., 2011). Analysis was restricted to periods when the mouse Gaussian smoothed velocity exceeded 1 cm/s. Spectrograms were calculated with a multitaper method with a 1 s sliding window. Coherence was calculated using a multitaper method over all run periods. Instantaneous frequency was defined by interpeak times of the band filtered trace during all run periods. LFP during run epochs was analyzed for 5 regions: CA1 pyr, sr, and slm; CA3 including pyr and sr; and DG including hilus and granule cell layers.

## QUANTIFICATION AND STATISTICAL ANALYSIS

Statistics were computed using custom software written in MATLAB (Mathworks). Figures 6 and S5–S6 were plotted with plotSpread function (MATLAB Central File Exchange). Statistical test used, exact n, and exact p value are in figure legends; test statistic values and degrees of freedom are in corresponding supplementary tables. In all cases, n represents number of animals. No data were excluded based on statistical tests. 3 mice were excluded from analysis due to poor viral expression. Sample sizes were based on previous studies (Gan et al., 2017; Lovett-Barron et al., 2014; Stefanelli et al., 2016; Xia et al., 2017). Central values plotted in Figures 2–5 and S2–S4 are means and individual points are mean per animal, as indicated in figure legends. Central values plotted in Figures 6 and S5 are linear mixed effects model (LMM) fixed effect coefficients β ± 95% confidence intervals and individual points are the fitted conditional response for each mouse, as indicated in figure legends.

Data for LMMs were drawn from events in the case of SWRs, time bins in the case of continuous measures (1 min for SWR rate, 100 ms for MUA, 1 s for all others), or peaks in the case of continuous frequency. Normality and independence of errors was confirmed visually; when errors were not Gaussian, they were always right skewed, and so a log transform was applied. We constructed an LMM using the ML method and the formula *feature ∼ group + (group|animal)* where group was vehicle or CNO for treatment comparison, PV-Cre or SST-Cre for vehicle baselines comparison, and male or female for sex comparison. We then used an F test to assess the evidence that the treatment fixed effect model was a better fit than the intercept-only model. To evaluate differences between treatment effects, we pooled data across two groups and used a likelihood ratio test to compare the model *feature ∼ treatment + (treatment|animal)* to the model *feature ∼ treatment:gentoype + (treatment:genotype|animal)* (or *feature ∼ treatment:sex + (treatment:sex|animal)* in the case of Table S6) and used a χ^2^ test to assess if the data were equally likely under both models. For table S1, we used paired t tests when data were normally distributed as shown by Shapiro-Wilk test and variances between groups were similar as shown by F test; we used Wilcoxon matched pairs signed rank tests otherwise. For correlations in Figures S2 and S3, data were normally distributed as shown by Shapiro-Wilk test, so we used Pearson correlations.

Significance threshold was set by the Holm-Bonferroni correction across an experiment with α = 0.05. Other family-wise error rate correction methods were tested with similar results. An experiment was defined as all comparisons made during either rest sessions (15 comparisons in Figures 2, 3, and S2) or run sessions (36 comparisons in Figures 4, 5, S3, and S4) in a single genotype or between genotypes. Only p-values that met this threshold are displayed as significant in figures and tables.

## DATA AND SOFTWARE AVAILABILITY

The data generated during this study will be deposited on CRCNS.org upon acceptance for publication. The custom software generated during this study is deposited at https://github.com/emilyasterjones/interneurons_modulate_drive.

